# Kinetic Explanations for the Sequence Biases Observed in the Nonenzymatic Copying of RNA Templates

**DOI:** 10.1101/2021.10.06.463393

**Authors:** Dian Ding, Lijun Zhou, Constantin Giurgiu, Jack W. Szostak

## Abstract

The identification of nonenzymatic pathways for nucleic acid replication is a key challenge in understanding the origin of life. We have previously shown that nonenzymatic RNA primer extension using 2-aminoimidazole (2AI) activated nucleotides occurs primarily through an imidazolium-bridged dinucleotide intermediate. The reactive nature and preorganized structure of the intermediate increase the efficiency of primer extension but remain insufficient to drive extensive copying of RNA templates containing all four canonical nucleotides. To understand the factors that limit RNA copying, we synthesized all ten 2AI-bridged dinucleotide intermediates and measured the kinetics of primer extension in a model. The affinities of the ten dinucleotides for the primer/template/helper complexes vary by over 7,000-fold, consistent with nearest neighbor energetic predictions. Surprisingly, the reaction rates at saturating intermediate concentrations still vary by over 15-fold, with the most weakly binding dinucleotides exhibiting a lower maximal reaction rate. Certain noncanonical nucleotides can decrease sequence dependent differences in affinity and primer extension rate, while monomers bridged to short oligonucleotides exhibit enhanced binding and reaction rates. We suggest that more uniform binding and reactivity of imidazolium-bridged intermediates may lead to the ability to copy arbitrary template sequences under prebiotically plausible conditions.

## INTRODUCTION

The absence of any known pathway for the efficient nonenzymatic copying of RNA templates that contain all four nucleotides is one of the main obstacles towards understanding the emergence of the first ribozymes and hence the origin of primordial RNA based life. Leslie Orgel pioneered the study of this subject by showing that imidazole and 2-methyl-imidazole activated 5′-monophosphate ribonucleotides could polymerize on RNA templates.(1–3) Recently, the mechanism of nonenzymatic primer extension mediated by 2-methyl-imidazolium-5′-phospho-ribonucleotides has been unraveled.(4) Surprisingly, the dominant reaction mechanism is not a simple S_N_2 reaction in which the 3′-OH of the primer attacks the 5′-phosphate of an adjacent template-bound activated mononucleotide. Instead, two activated monomers first react with each other to form a covalent intermediate in which the two mononucleotides flank a central imidazolium group (Figure 1A). The intermediate dinucleotide binds to the template at the +1 and +2 positions via Watson-Crick base pairs, after which the 3′-hydroxyl group of the primer attacks the proximal phosphate group, resulting in a +1-extension product. The nucleotide bound at the +2 position of the template, together with the imidazolium-phosphate moiety, acts as the leaving group in this reaction. Primer extension via direct attack on an activated monomer can occasionally occur but this reaction is far slower than the reaction with an imidazolium-bridged dinucleotide.(5) Sequencing of the products of primer extension has also shown that monomer-mediated primer extension is highly error prone compared to primer extension with imidazolium-bridged dinucleotides.(6, 7)

**Figure 1:**
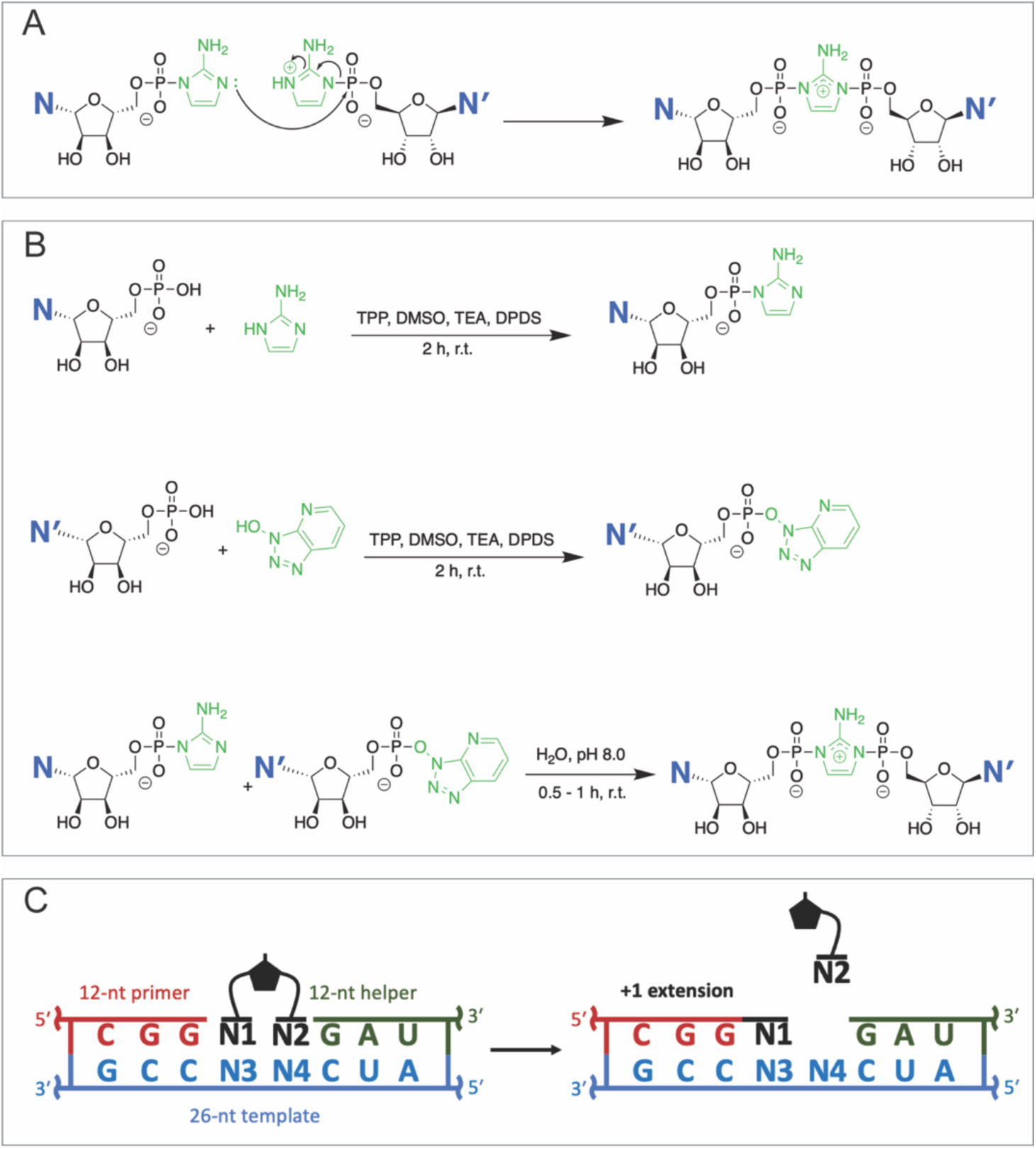
(A) The spontaneous formation of 2-aminoimidazolium-bridged dinucleotides in solution. (B) Scheme for chemical synthesis of imidazolium-bridged dinucleotides. (C) Scheme for primer extension experiments. Primer extension reactions were carried out with 1.5 µM primer, 2.5 µM template, 3.5 µM helper, 100 mM MgCl_2_ and 200 mM Tris pH 8.0.

Nucleotide activation with 2-aminoimidazole leads to increased rates and yields of primer extension compared to 2-methylimidazole or imidazole.(8) The superior properties of 2-aminoimidazole are attributable at least in part to the greater stability of the corresponding bridged dinucleotide intermediates, which accumulate to higher levels resulting in more efficient primer extension. Moreover, a potentially prebiotic pathway for the synthesis of 2-aminoimidazole shares a common mechanism with and can occur in the same reaction mixture as that of 2-aminooxazole, a precursor of prebiotic nucleotide synthesis(9). For these reasons we believe that 2-aminoimidazole activated ribonucleotides are the best available model for the study of nonenzymatic template-directed primer extension reactions.

Despite the enhanced rate and fidelity of primer extension via imidazolium-bridged dinucleotides, this process is only efficient on G+C templates, while templates containing A, G, C and U are copied very poorly. Although some bias against A and U is to be expected given the weaker A:U base-pair compared to the G:C base-pair, a deeper mechanistic understanding of the template copying process is required to solve the problem of copying mixed sequence templates. Until now, a detailed kinetic study of primer extension with 2-amino-imidazolium-bridged dinucleotides has only been done with the dicytidine intermediate (C*C).(10) This study provided a kinetic framework for the synthesis and breakdown of the intermediate, as well as the greater reactivity of the dinucleotide intermediate relative to the activated monomer. However, since only C*C was studied, this study did not address the problem of copying arbitrary RNA sequences. Recently, our lab has reported the use of deep sequencing to study nonenzymatic RNA primer extension on random sequence templates(7). As expected, we observed an initial bias favoring primer extension with G and C, but surprisingly continued primer extension became even more strongly biased in favor of C. Template sequencing suggested that this bias resulted from the superior activity of the C*C dinucleotide relative to all other N*N dinucleotide intermediates. In order to understand the kinetic basis for this result, kinetic studies of the other N*N intermediates are needed. Here we report the chemical synthesis and characterization of all ten dinucleotide intermediates (A*A, C*C, G*G, U*U, A*C, A*G, A*U, C*G, C*U and G*U) and the concentration dependent reaction rates of each dinucleotide when bound to its respective Watson-Crick base pairing template. Our results reveal a surprisingly broad range of affinities and reaction rates. Intriguingly, the identity of the downstream nucleotide significantly affects the rate of primer extension. NMR studies of sugar pucker conformation provide a potential explanation for the preferential incorporation of C during RNA copying. Preliminary kinetic studies of bridged dinucleotides containing the noncanonical nucleotide 2-thio-U, as well as monomers bridged to short oligonucleotides, suggest strategies for the further optimization of the nonenzymatic copying of arbitrary RNA sequences.

## MATERIAL AND METHODS

### Synthesis and characterization of 2-aminoimidazolium-bridged dinucleotide intermediates

The homo-dimers A*A, C*C and U*U were synthesized by mixing 1 equivalent of nucleoside 5′-monophosphate with 0.45 equivalent of 2-aminoimidazole (2AI) in dry dimethyl sulfoxide (DMSO), then adding 20 equivalents of triethylamine (TEA), 10 equivalents of triphenylphosphine (TPP) and 10 equivalents of 2,2′-dipyridyldisulfide (DPDS). After incubation for 30 mins, the product was precipitated in acetone with sodium perchlorate. The precipitant was washed twice by mixture of acetone and diethyl ether (1:1, v/v) and dried under vacuum. Then the dry pellet was resuspended in 20 mM triethylamine-bicarbonate buffer (TEAB), pH 8.0, and purified by reverse-phase flash chromatography, with a 50 g C18Aq column. The desired products were separated from other compounds by reverse phase flash chromatography over 12 CVs of 0–10% acetonitrile in 2 mM TEAB buffer (pH 8.0). We did not use this method for G*G, because the low solubility issue of guanosine 5′-monophosphate (GMP) prevented us from mixing GMP with 2AI at the desired ratio.

To synthesize G*G and the six hetero-dimers (N*N′), we first synthesized and purified 2AI activated N monomer and 1-hydroxy-7-azabenzotriazole (HOAt) activated N′ monomer, as described above except that 10 equivalents of 2AI or HOAt were added. To overcome the poor solubility of GMP in DMSO, GMP was first dissolved in water together with 10 equivalents of 2AI or HOAt and 100 mM TEA, flash frozen and lyophilized, and then dissolved in DMSO for further reactions. Then purified 2AI-activated N and HOAt-activated N′ were mixed in a 1:1 molar ratio in water, at pH 8.0 (adjusted by NaOH/HCl). HOAt is a good leaving group but cannot form a 5′-5′ bridged intermediate like 2AI. After incubation for 30–60 minutes, the 2AI intermediate dimer N*N′ was the predominant product, alongside byproducts such as N*N, N′*N′, and activated monomers. The desired product was then purified by preparative scale HPLC using a C18 reverse phase column and a solvent gradient (A) 2 mM aqueous TEAB buffer, pH 8.0, (B) acetonitrile. The gradients for each different bridged dinucleotide are described in the supplementary information.

The synthesis of the noncanonical bridged dinucleotides sU*sU and sC*sC started with phosphorylation of 2-thiouridine and 2-thiocytidine to form the 5′-monophosphate sUMP and sCMP. 1 equivalent of the 5′-hydroxyl nucleotide was dissolved in an ice-chilled solution of 80 equivalents of trimethyl phosphate, 3.6 equivalents of phosphoryl chloride, and 0.4 equivalent of water. Then four separate portions of *N,N*-diisopropylethylamine (0.52 equiv. each) were added dropwise to the stirred solution with 20 min intervals at 0°C. The reaction mixture was stirred for another 20 min and quenched with 1M TEAB, pH 7.6. The mixture was purified first by reverse phase flash chromatography over 5 CVs using 25 mM TEAB buffer (pH 7.5). Fractions containing product were lyophilized and the resultant syrup was purified again by reverse phase chromatography over 10 CVs of 0–10% acetonitrile in 25 mM TEAB buffer (pH 7.5) to give sUMP or sCMP. The activation of sUMP and sCMP using 2AI led to a low yield (10 ∼ 20%) and we were unable to synthesize HOAt-activated sU or sC. Therefore, sU*sU and sC*sC were synthesized by activating the nucleotide monophosphate using 2AI-activated monomer as follows. sUMP or sCMP was activated using 2AI as described above to form 2AI-sU or 2AI-sC. Then 1 equivalent of 2AI-sU or 2AI-sC and 1.15 equivalent of sUMP or sCMP were incubated with 30 equivalents of TPP and 30 equivalent of DPDS for 2h in DMSO using a similar activation procedure. The desired product was then purified by reverse phase flash chromatography over 25 CVs of 0–10% acetonitrile in 2 mM TEAB buffer (pH 7.5). Monomer-bridged-oligonucleotides were made from 2AI activated oligonucleotides and HOAt-activated-pA. The 5′-phosphorylated oligonucleotides (pCG, pCGC, and pCGCA) were prepared by solid-phase RNA synthesis. The oligonucleotide 5′-monophosphate was then activated using 40 equivalents of 2AI, 40 equivalents of TPP, and 40 equivalents of DPDS in DMSO and TEA for 6 hours following a similar activation procedure as described above. The 2AI-activated-oligonucleotide was then mixed with 1.5 equivalent of HOAt-activated-pA in water at pH 8.0 for 1h. The mixture was purified by preparative scale HPLC using a C18 reverse phase column over 27 CVs of 0–10% acetonitrile in 25 mM TEAB buffer (pH 7.5).

The detailed synthesis procedure for each individual bridged intermediate as well as their characterizations are included in the supplementary data.

### Nonenzymatic primer extension reactions

Primer/template/helper duplexes were prepared in an annealing buffer at 5x the final concentration. Solutions containing 7.5 µM primer, 12.5 µM template, 17.5 µM helper, 50 mM Tris-Cl (pH 8.0), 50 mM NaCl and 1 mM EDTA (pH 8.0) were heated at 95°C for 30 s and then slowly cooled to 25°C at a rate of 0.1°C/s in a thermal cycler machine. The annealed products were then diluted five-fold into the primer extension reaction buffer containing 200 mM Tris-Cl, pH 8.0, and 100 mM MgCl_2_. Stock solutions of imidazolium-bridged dinucleotide intermediates were freshly prepared and added to the buffer with annealed primer/template/helper duplexes to initiate primer extension reactions. At each time point, 0.5 µL of reaction sample was added to 25 µL quenching buffer containing 20 mM EDTA, 1x TBE and 10 µM of an RNA complementary to the template.

### Urea-PAGE analysis

Primer extension products were resolved by 20% (19:1) polyacrylamide gel electrophoresis (PAGE) with 7 M urea, in 1x TBE gel running buffer. The gels were scanned with an Amersham Typhoon RGB Biomolecular Imager (GE Healthcare Life Sciences). The fluorescently-labeled primer and extended primer bands were visualized, and then quantified using ImageQuant TL software to obtain relative band intensities.

### NMR measurement of sugar conformations

Proton NMR spectra of all imidazolium-bridged dinucleotides were acquired on a Varian Oxford AS-400 NMR spectrometer (for details see the Supplementary Information). Proton NMR peak assignments were performed with the help of coupling constants and 2D COSY NMR. For all the hetero-dinucleotide intermediates, the two different nucleotides in the molecule have well separated H1ʹ hydrogen peaks. The ^3^J_H1ʹ-H2ʹ_ coupling constant was measured from the NMR peak corresponding to H1’ only, because the H2’ peak overlaps with other peaks for most of the bridged dinucleotides. The ratio of the fraction of 3’-endo to 2’-endo sugar pucker conformation was calculated based on the equation: C3’-endo/2’-endo = J/(10-J) where J = ^3^J_H1ʹ-H2ʹ_.(11)

## RESULTS

### Stability of imidazolium-bridged dinucleotides

To investigate the reasons for the strong sequence bias in nonenzymatic RNA copying, we conducted a comparative study of the kinetics of template copying using purified imidazolium-bridged dinucleotide intermediates. In order to avoid artifacts due to possible differences in chemical stability, we first measured the rates of hydrolysis of selected N*N intermediates. The degradation rate of the homo-dinucleotide C*C has previously been reported as 0.17 h^-1^ (half-life ∼245 min)(10), under slightly different buffer conditions than used here (90 mM Tris (pH 8.3-8.4) and 90 mM MgCl_2_). We used proton NMR to follow the degradation of A*C and A*U, two representative hetero-dinucleotides, under our primer extension reaction conditions (200 mM Tris 8.0 and 100 mM MgCl_2_, Figure S1). We found that both A*C and A*U degraded by hydrolysis to give 2AI-activated monomers and unactivated 5′-monophosphate monomers; these compounds subsequently reacted to form imidazolium-bridged homo-dinucleotides and other products. As a result, the overall degradation process did not follow simple pseudo-first-order reaction kinetics. Nevertheless, from the time-course curves, half-lives for A*C and A*U of ∼135 min and ∼120 min respectively were estimated (Figure S1). At 10 min, 95% and 94% of A*C and A*U respectively remained unhydrolyzed. We conclude that primer extension reaction rates obtained within ten minutes should be reasonably accurate and should allow for valid comparisons between different imidazolium-bridged dinucleotides.

### Nonenzymatic primer extension reactions with 5 mM intermediate

For an initial comparison of the reaction rates of the ten different N*N intermediates, we set up ten different primer extension reactions, each with 5 mM of one intermediate and its complementary template. The experimental design is illustrated in Figure 1C, with the 26-nt template, 12-nt primer and 12-nt downstream helper (5′-OH). Each intermediate N1*N2 binds to the template through two Watson-Crick base-pairs with the nucleotides N3 and N4, and is sandwiched between the 3′-guanosine of the primer and the 5′-guanosine of the downstream helper oligonucleotide. Without the stabilizing effect of the stacking interaction with the downstream helper, the reactions of some intermediates (e.g., U*U) are barely detectable. This experimental design allowed us to make a direct comparison of the pseudo-first-order reaction rates (*k*_obs_) of all ten intermediates in sixteen parallel sets of reactions, as shown in Figure 2.

**Figure 2:**
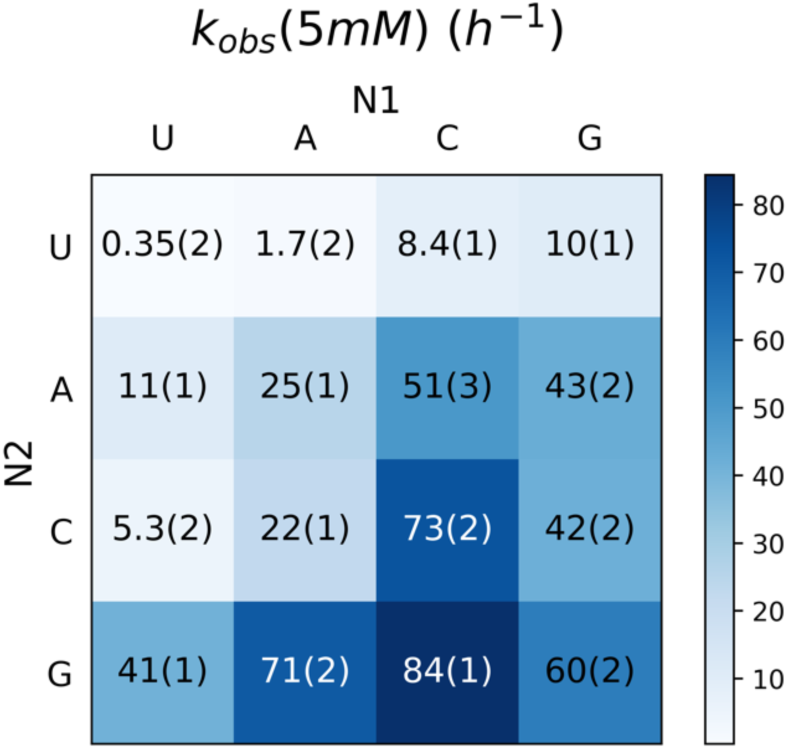
*k*_obs_ of nonenzymatic primer extension reactions of imidazolium-bridged dinucleotide intermediates N1*N2 at 5 mM. Note that intermediates N1*N2 and N2*N1 are identical, but are bound to templates with reverse orientation e.g. 5′-N3-N4-3′ and 5′-N4-N3-3′. The experimental design is described in Fig 1C.

At a concentration of 5 mM for each imidazolium-bridged dinucleotide, the observed rates of primer extension varied over a range of 240-fold, with the lowest *k*_obs_ of 0.35 h^-1^ for U*U and the highest *k*_obs_ being 84 h^-1^ for C*G. The reaction rates in Figure 2 are consistent with previous studies of 2-methylimidazole activated mono-nucleotides, which form analogous imidazolium-bridged homo-dinucleotide intermediates, in that primer extension with G*G and C*C are the most rapid, while primer extension with A*A is slower(4). However, when we examined the reaction rates of the hetero-dinucleotide intermediates, we found surprising differences depending on the 5′-3′ orientation of the intermediate on the template. For example, primer extension with 5′-U*G (41 h^-1^) is four-fold faster than with 5′-G*U (10 h^-1^) even though this intermediate has the same Watson-Crick base pairing in both cases. Similarly, primer extension with 5′-U*A (11 h^-1^) is six-fold faster than with 5′-A*U (1.7 h^-1^). Comparing the reaction rates in the columns of Figure 2 shows that the identity of the downstream base significantly affects the primer extension rate. With the same N1, the trend for *k*_obs_ generally follows N1\***U** < N1\***C** ≤ N1\***A** < N1\***G**, consistent with a favorable influence of the stronger C•G vs. A•U base-pair, combined with the stronger stacking interaction of purines (A, G) than pyrimidines (C, U). For a given N2, the trend of *k*_obs_ is **U**\*N2 < **A**\*N2 < **G**\*N2 ≤ **C**\*N2 in general, consistent with previous studies showing that primer extension rates with C and G are robust, following by A, while the extension of U is very poor. However, when N2=G, the trend of N1 reaction rate is U < G < A < C. Although the extension rates of all four nucleobases are rapid, this different order was surprising, and led us to ask whether the different observed rates reflected differences in binding or reactivity for the different intermediate/template combinations.

### Kinetic study of nonenzymatic primer extension reactions

We sought to determine whether the differences in the observed primer extension reaction rates were due to differences in the affinities or the reactivities of the imidazolium-bridged dinucleotide intermediates. To answer this question, we measured the rates of primer extension as a function of the concentration of each intermediate. This allowed us to obtain the observed maximal rate (*k*_obs max_), when the template is saturated with the intermediate (Figure 3). We also fitted the data to Michaelis-Menten equation and generated values for K_m_, which reflects the affinity of each dimer (Figure 4, S2). Though Michaelis-Menten models are used for enzyme kinetic studies, they are also suitable for the case of nonenzymatic primer extension reactions. Firstly, the concentrations of imidazolium-bridged dinucleotides (“substrates”) are three orders of magnitudes higher than the concentrations of primer/template/helper duplexes (“enzymes”). Secondly, the reactions are also two step reactions, in which a rapid reversible step of dinucleotide binding is followed by an irreversible step of phosphodiester bond formation catalyzed by the primer/template/helper complex. As is the case with enzymatic reactions, K_m_ is not necessarily equal to K_d_, but is a close approximation of K_d_ as long as the on and off rates of the ‘substrate’ imidazolium-bridged dinucleotide are fast relative to the rate of the chemical step. Due to the reactive nature of the dinucleotide substrates, we have not yet been able to directly measure K_d_ values by physical methods. To avoid the effect of differing affinities, we compared the reactivity of each dimer under conditions of template saturation (*k*_obs max_). The results showed a 15-fold difference between the lowest (5.7 h^-1^ for U*U) and the highest reaction rate (85 h^-1^ for C*G). Under saturation conditions the variation in reaction rates is decreased relative to the scenario in which all intermediates were present at the same low concentration, consistent with significant differences in affinity contributing to the larger range of reaction rates when all intermediates were present at the same concentration. Surprisingly, however, the trends in reaction rates were similar at 5 mM and at saturating concentrations. For a given N1, reaction rates followed N1*U < N1*A < N1*C < N1*G for *k*_obs max_. We conclude that different imidazolium-bridged dinucleotides have intrinsic differences in reactivity.

**Figure 3:**
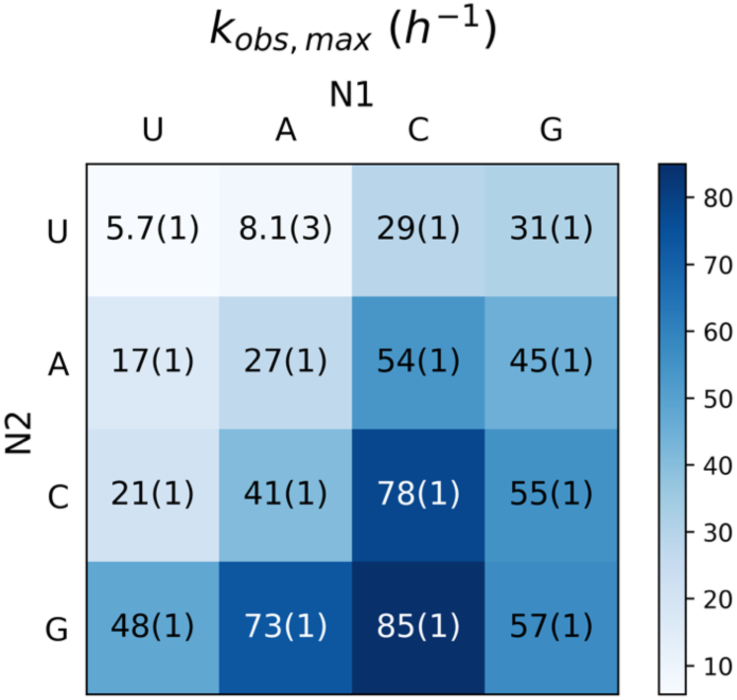
*k*_obs max_ of nonenzymatic primer extension reactions from 16 Watson-Crick base pairing of dinucleotide intermediates N1*N2. The experimental design is as described in Fig 1C.

**Figure 4:**
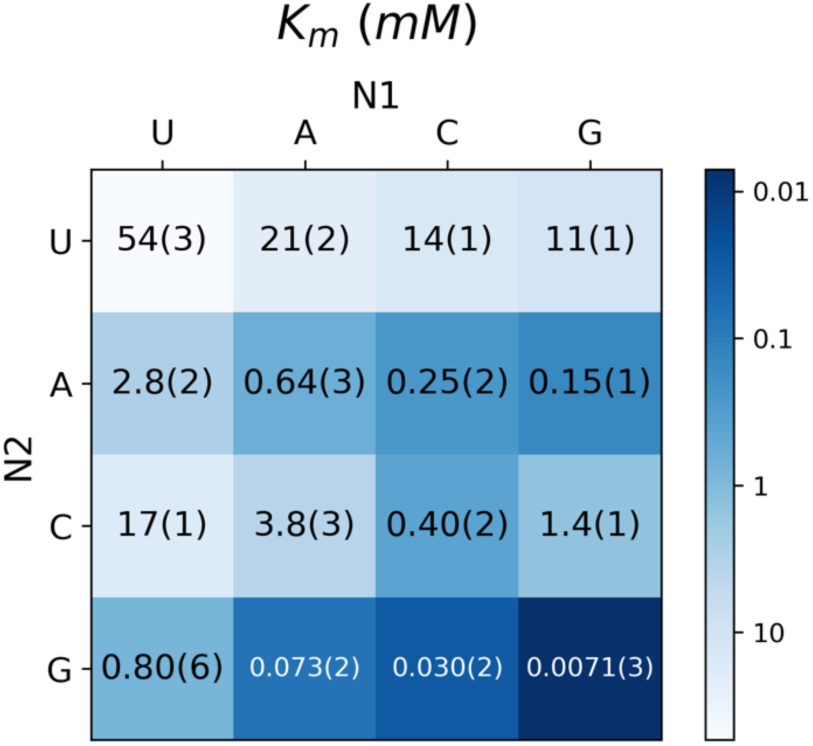
K_m_ of imidazolium-bridged dinucleotide intermediates in nonenzymatic primer extension reactions, for all 16 possible Watson-Crick N1*N2/template combinations. The experimental design is as described in Fig 1C.

Next, we measured the affinities of each imidazolium-bridged dinucleotide and asked what factors affect template binding. Figure 4 shows that the highest K_m_ is 54 mM (U*U), while the lowest K_m_ is 0.0071 mM (G*G); the difference is almost four orders of magnitude. For a given N1, the trend is N1*U > N1*C > N1*A > N1*G. For the same N2, K_m_ the trend is U*N2 > A*N2 > C*N2 > G*N2 in general. The K_m_ of G*C is surprisingly high at 1.4 mM (vs. 0.03 mM for C*G), indicating relatively weak binding affinity. This large difference must reflect that importance of stacking interactions in stabilizing the binding of imidazolium-bridged dinucleotides. This is also reflected in the 14-fold difference in affinities between G*U (K_m_ = 11 mM) and U*G (K_m_ = 0.80 mM). Considering all pyrimidine*purine intermediates, the affinity is always greater when the pyrimidine is upstream. That is, the K_m_ of U*A is less than the K_m_ of A*U, the K_m_ of C*A is less than the K_m_ of A*C, and the K_m_ of C*G is less than the K_m_ of G*C. The differences vary from 8 to 47 fold. The effect of binding orientation is minor in the case of purine*purine and pyrimidine*pyrimidine imidazolium-bridged intermediates, e.g. the K_m_ of U*C is only 1.2-fold higher than the K_m_ of C*U, and the K_m_ G*A is only 2-fold higher than the K_m_ of A*G. These effects are likely due to the sum of the stacking interactions with the guanosines that flank the bound imidazolium-bridged dinucleotides, e.g. the stacking interactions 5′-*GN*_*1*_-3′ and 5′-*N*_*2*_*G*-3′. It has been previously reported that the 5′-*Pu-Py*-3′ orientation results in greater energy of stacking than the 5′-*Py-Pu*-3′ orientation.(12) As a result, when N_1_ is a pyrimidine and N_2_ is a purine, the sum of the stacking interactions is predicted to be stronger than for the reverse case.

As noted above there is a correlation between the reactivities and the affinities of the imidazolium-bridged dinucleotide intermediates. One possible explanation is that strong binding of the intermediate stabilizes the configuration that is optimal for nonenzymatic primer extension. For example there could be situations where one nucleotide of the dinucleotide intermediate is bound, but the other one is fluctuating between the bound and the unbound state. If N1 of the dinucleotide is only Watson-Crick paired to the template a fraction of the time that the dinucleotide is bound, the rate of the primer extension reaction would be correspondingly decreased. More detailed studies of the physical characteristics of the bound intermediates are required to further understand their behavior in primer extension.

### The effect of ribose configuration on the reaction

The main exception to the trends in binding and reactivity discussed above is the anomalously high reactivity of the imidazolium-bridged dinucleotides in which N1 is C. This observation is consistent with the results from deep sequencing of the products of nonenzymatic primer extension on random templates, in which primer extension is dominated by the incorporation of C.(6) This cannot be explained solely by affinity, since the binding of G should generally be stronger. What then could account for the greater reactivity if imidazolium-bridged dinucleotides with C at the N1 position?

We asked if the conformational preference of the ribofuranose rings of the bridged dinucleotides could potentially explain this anomalous reactivity. The furanose rings of ribonucleotides can pucker in two different ways: the C3ʹ-endo and the C2ʹ-endo conformations. Although ribonucleotides generally prefer to adopt the C3ʹ-endo configuration, the two conformations are close in energy and can easily interconvert in solution. Therefore, the conformational behavior of most nucleotides can be modeled as a two-state equilibrium; this has been extensively characterized by NMR.(13) Previous studies suggest that the C3ʹ-endo conformation for the template and 3ʹ-terminal primer position is optimal in primer extension reactions.(14–16)

Activated 5′-GMP has been shown to undergo a sugar pucker switch from predominantly C2ʹ-endo in solution to C3ʹ-endo upon binding to an RNA template.(17) In addition, crystallographic studies show that the G*G intermediate exhibits only the C3′-endo configuration when bound to the template downstream of a primer.(15)

To understand the factors that influence the rates of nonenzymatic primer extension reactions, we determined the sugar pucker conformation of each ribonucleotide at the N1 position of all 10 imidazolium-bridged dinucleotides. We determined the ratio between the C3′- and C2′-endo conformations by NMR, making use of the well-established correlation between the ^3^J coupling constant between the H1ʹ and H2ʹ atoms (^3^J_H1ʹ-H2ʹ_) (13, 17), and the furanose ring pucker (Figure 5, right). A lower value of ^3^J_H1ʹ-H2ʹ_ indicates a higher proportion of the 3′-endo configuration. Our observations show that cytidine exhibits a higher degree of 3′-endo configuration relative to other nucleotides at the N1 position. We hypothesized that this higher fraction of 3′-endo conformation could potentially contribute to the faster reaction rate of imidazolium-bridged dinucleotides with C at the N1 position.

**Figure 5:**
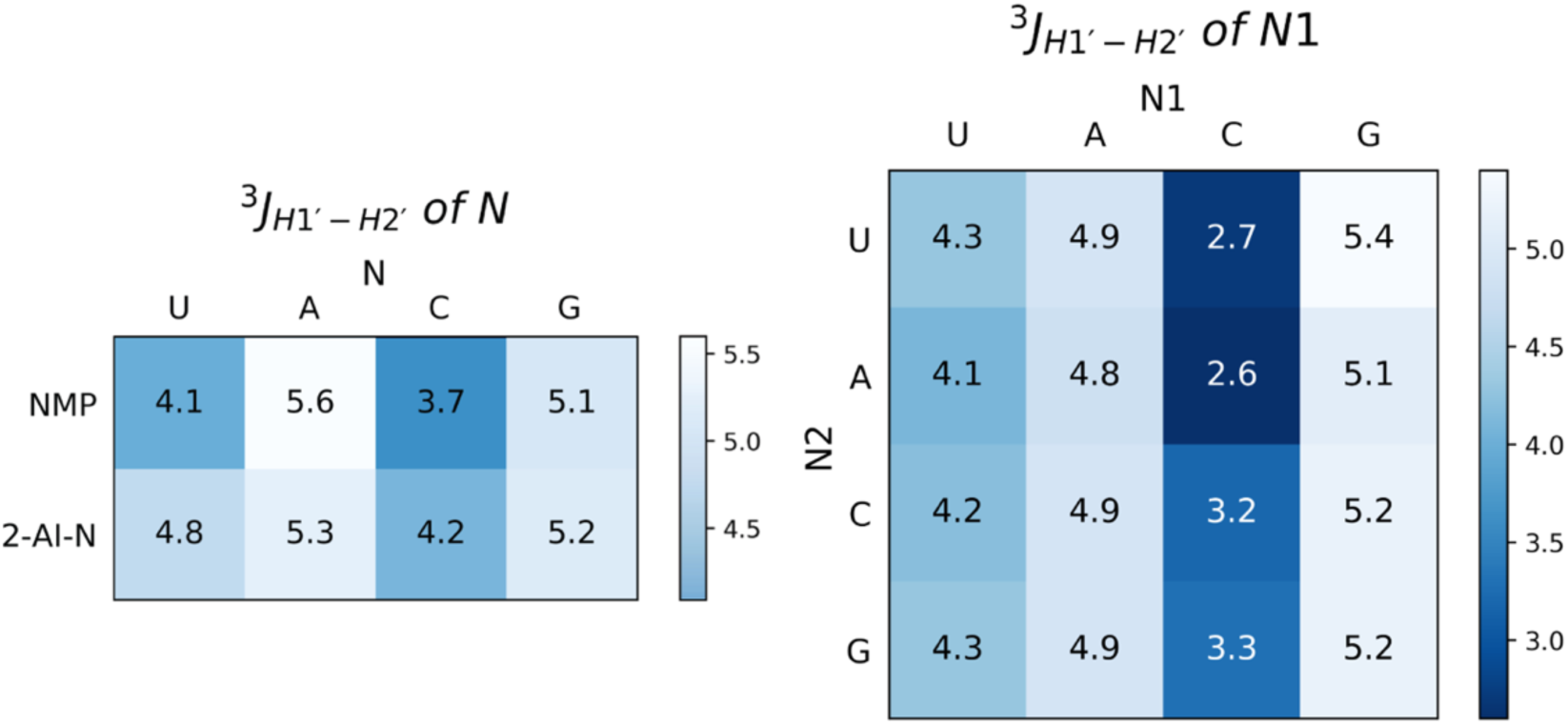
Left: ^3^J_H1ʹ-H2ʹ_ of the nucleotide in monophosphate form and in the activated monomer form measured at pH 8.0. Right: ^3^J_H1ʹ-H2ʹ_ of N_1_ nucleotide in the intermediate N_1_*N_2_ measured at pH 8.0. A lower value of ^3^J_H1ʹ-H2ʹ_ represents a higher degree of 3′-endo configuration.

To further test the effect of the ribose sugar pucker of the substrate of primer extension reactions, we also investigated the sugar pucker of ribonucleotide monomers. We measured the values of ^3^J_H1ʹ-H2ʹ_ of the four 5′-monophosphate ribonucleotides, with and without 2-aminoimidazolium activation, in solution (Figure 5, left). The ^3^J_H1ʹ-H2ʹ_ values for NMPs has been reported previously with minor differences from this work.(18) Cytidine and uridine show stronger preferences for the C3ʹ-endo conformational compared to guanosine and adenosine. To determine whether the 3′-endo conformation leads to a higher reaction rate for activated monomers, we conducted primer extension reactions with only one nucleotide binding site on the template (Figure S3). Here, 2AI-activated ribonucleotide monomer (*N) is stacked with G at 3′-end of primer and with another G at the 5′-end of the downstream helper oligonucleotide. Although the affinities of *C and *U are much lower than *G and *A (as expected due to decreased staking energies), *C has higher *k*_obs max_ compared to *G, and quite surprisingly *U has higher *k*_obs max_ compared to *A.

To further examine the effect of sugar pucker on reactivity and to control for any N1 or N2 nucleobase-identity effects, we synthesized the imidazolium-bridged dinucleotides of four sugar-modified guanosines with different conformational preferences, and measured their performance in the primer extension reaction without the downstream helper (Figure 6). By using the same guanosine base in these modified dinucleotides, any difference in the primer extension behavior should be a result of the different 2ʹ functional group on the sugar. The maximal rate of reaction correlates strongly with the ratio of the C3ʹ-endo to C2ʹ-endo conformers, as calculated from the values of ^3^J_H1ʹ-H2ʹ_ (Figure 6C). The fastest reacting imidazolium dimer is the locked-nucleic acid (LNA), which can only exist in the C3ʹ-endo conformation; the slowest reacting dinucleotide is the DNA version, which prefers the C2ʹ-endo conformation. Our results confirm that the primer extension reaction is faster when the imidazolium-bridged intermediate is pre-organized with the nucleotide sugars in the C3ʹ-endo conformation.

**Figure 6:**
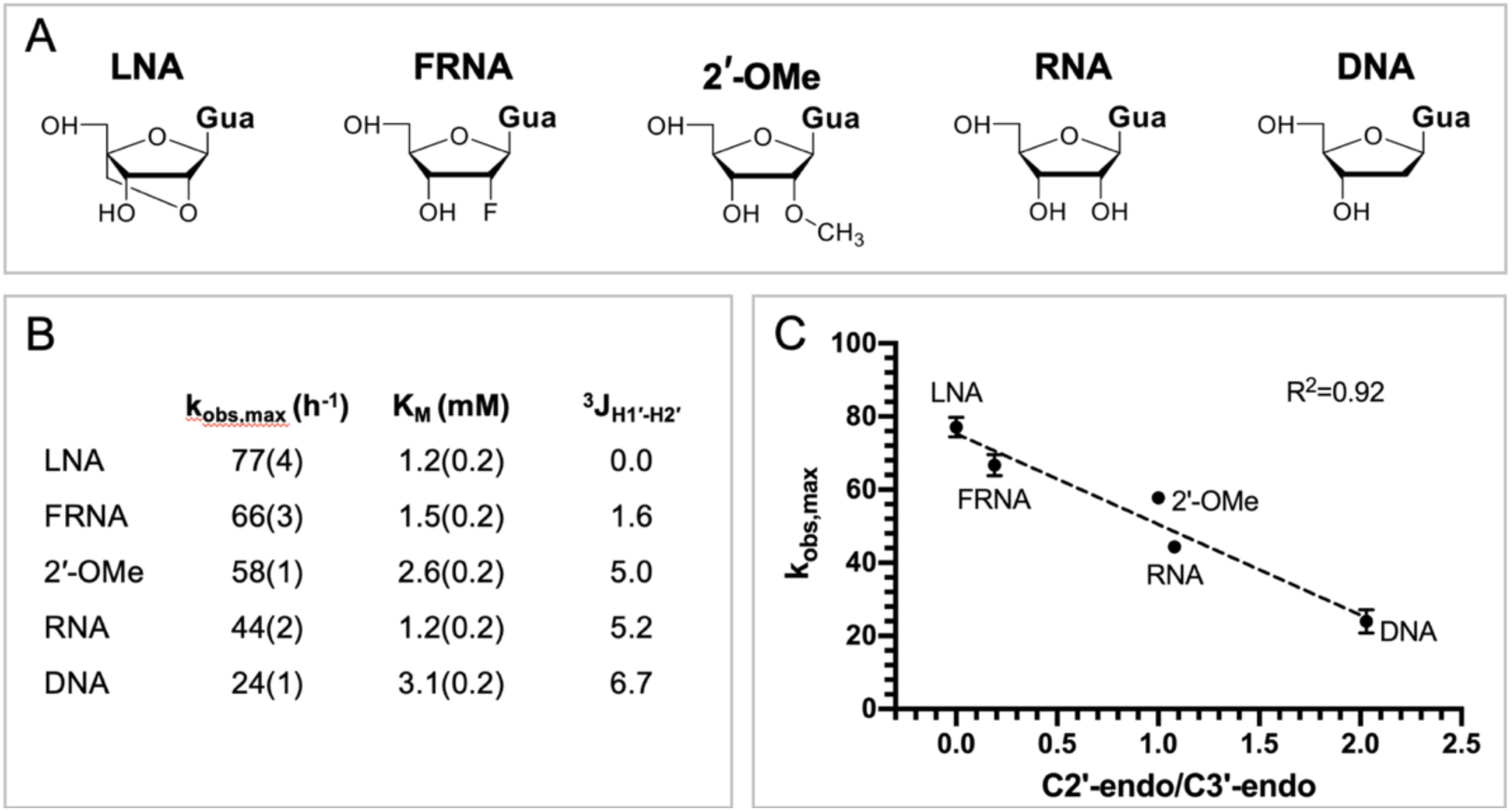
Primer extension with 2ʹ modified di-guanosine imidazolium-bridged dinucleotides. (A) Chemical structures of different modified guanosine nucleotides used to form the dinucleotide intermediates. (B) Kinetic parameters and ^3^J_H1ʹ-H2ʹ_ for each modified di-guanosine imidazolium-bridged dinucleotide (C) The correlation between the maximal rate of the reaction and the ratio of the two conformers.

### Potential solutions to the weak binding and poor reactivity of specific dinucleotide intermediates

We propose two strategies to overcome the poor efficiency of primer extension that results from the poor binding and reactivity of certain imidazolium-bridged dinucleotides. First, we suggest that monomers that are imidazolium-bridged to oligonucleotides will bind more strongly to the template, and may therefore enhance the rate of nonenzymatic primer extension. Our lab has previously demonstrated that an activated helper oligonucleotide catalyzes nonenzymatic primer extension with an activated monomer that is sandwiched between the primer and the helper oligonucleotide.(19) With the recent discovery of the imidazolium-bridged dinucleotide intermediate, it now seems likely that the catalytic effect of the activated downstream oligonucleotide is due to the formation of monomer-bridged-oligonucleotides. In order to test this hypothesis directly, we synthesized a series of monomer-bridged-oligonucleotides and measured their rates of nonenzymatic primer extension as a function of concentration (Figure 7). For this set of experiments, we used a primer with a 3ʹ-terminal C and no downstream helper oligonucleotide, to minimize the effect of stacking interactions on binding. Remarkably, compared to A*C, A*CG is 7-fold better in maximum nonenzymatic primer extension rate, over 20-fold better in binding, and almost 150-fold better in *k*_obs max_ / K_m_. As the bridged oligo gets longer, the *k*_obs max_ reaches a plateau; this *k*_obs max_ plateau is comparable to the highest *k*_obs max_ observed in the sandwich model with bridged dinucleotides. This finding suggests that the large differences observed between the affinities of different bridged dinucleotides can be reduced with the use of monomer-bridged-oligos. Moreover, since the affinity increases significantly as the intermediate gets longer, the maximum reaction rate is reached at much lower concentrations (Figure 7B).

**Figure 7:**
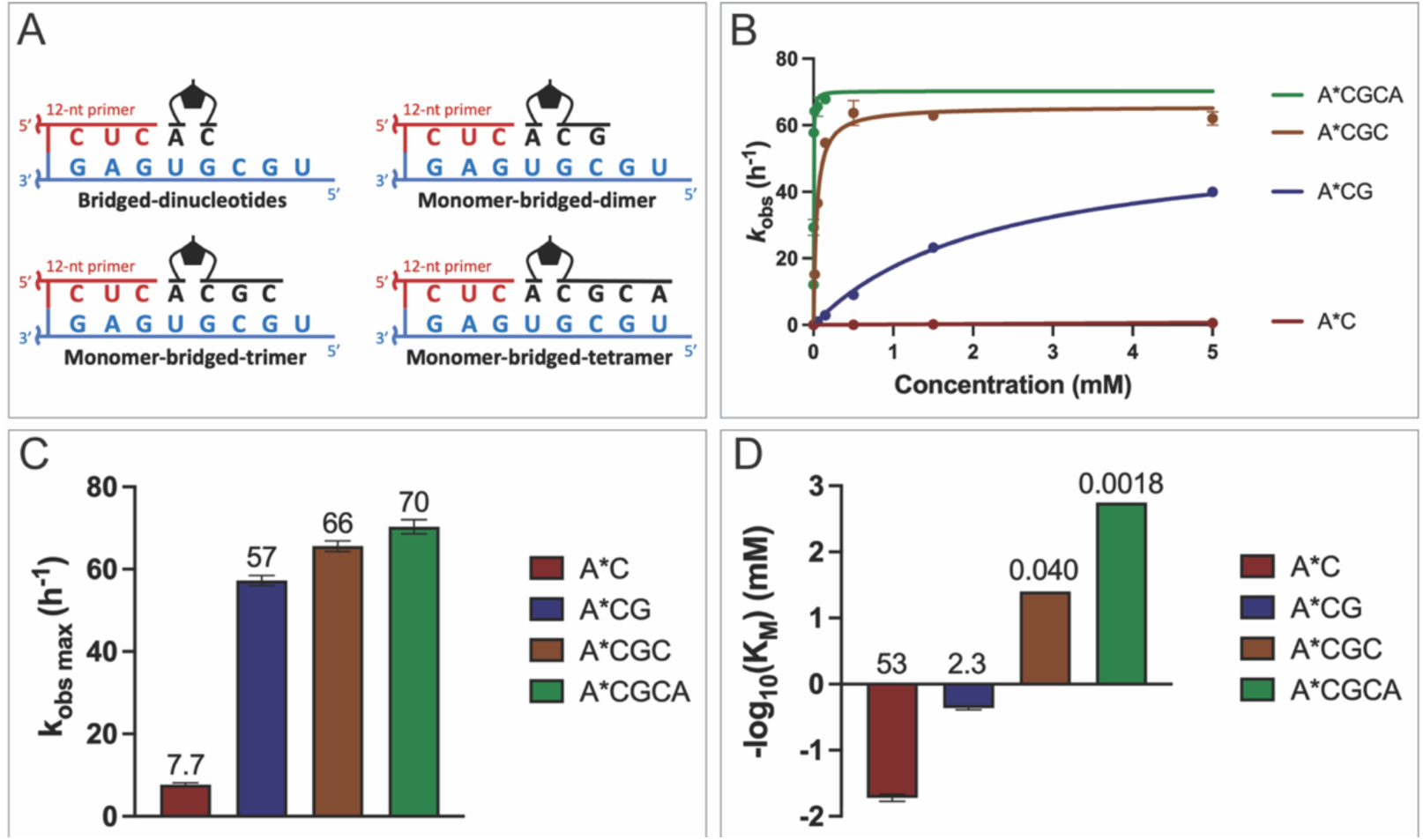
Primer extension with monomers imidazolium-bridged to oligonucleotides. The primer ends in a 3ʹ-terminal C; there is no downstream helper. (A) Schematic representation of the nonenzymatic primer extension setup. (B) Michaelis-Menten curves plotted for 0-5 mM of the monomer-bridged-oligonucleotide intermediates. See the SI for the Michaelis-Menten plots of each individual monomer-bridged oligonucleotide. The comparison of (C) *k*_obs max_ and (D) -log_10_(K_M_) between A*C, A*CG, A*CGC, A*CGCA are shown. The values of *k*_obs max_ and K_M_ are indicated above each bar.

As an alternative potential solution to the problems resulting from the wide range of affinities of the different bridged dinucleotides, we propose that the use of certain noncanonical nucleotides may lead to more uniform binding and thus more uniform rates of nonenzymatic primer extension. The stereoelectronic character of the nucleobase can tune the sugar pucker confirmation of a nucleotide,(20) and 2-thiolated pyrimidines (2-thioU, sU; 2-thio-C, sC) have been shown to have an increased preference for the C3ʹ-endo configuration.(21) Furthermore, the sU:A base pair is stronger than U:A base pair,(22) while the sC:I (inosine) base pair(23) is weaker than the C:G base pair because it has 2 instead of 3 hydrogen bonds per base pair. To examine the conformations, affinities and rates of the corresponding imidazolium-bridged dinucleotides, we synthesized the 2-thio-uridine homodimer (sU*sU) and the 2-thio-cytidine homodimer (sC*sC). The ^3^J_H1ʹ-H2ʹ_ value of sU*sU is around 2.3, lower than that of any other nucleotide reported in Figure 5. Surprisingly, sC*sC has a ^3^J_H1ʹ-H2ʹ_ around 0, indicating an almost entirely C3ʹ-endo configuration in solution. We then measured the nonenzymatic primer extension rate of the two homo-dimers in the sandwich model with sU:A and sC:I base pairs at the binding pocket (Figure 8). The sU*sU dinucleotide shows almost 8-fold increase and the sC*sC has almost 3-fold increase in the *k*_obs max_. Note that the sC*sC has an extremely high *k*_obs max_ = 220 h^-1^, which means almost half of the primer can be extended in just 8 seconds. In addition, the Km of sU*sU has been improved by about 2-fold, while the binding of sC*sC on a 3ʹ-II template is 20-fold weaker compared to C*C on a 3ʹ-GG template. (Figure S2, S6). As a result, the over 130-fold affinity difference between the two bridged dinucleotides has been decreased to only 3-fold. We suggest that the use of noncanonical nucleotides and monomer-bridged-oligonucleotides may allow for significantly increased rates of nonenzymatic primer extension and decreased differences between the affinities of different bridged dinucleotides for mixed sequence templates.

**Figure 8:**
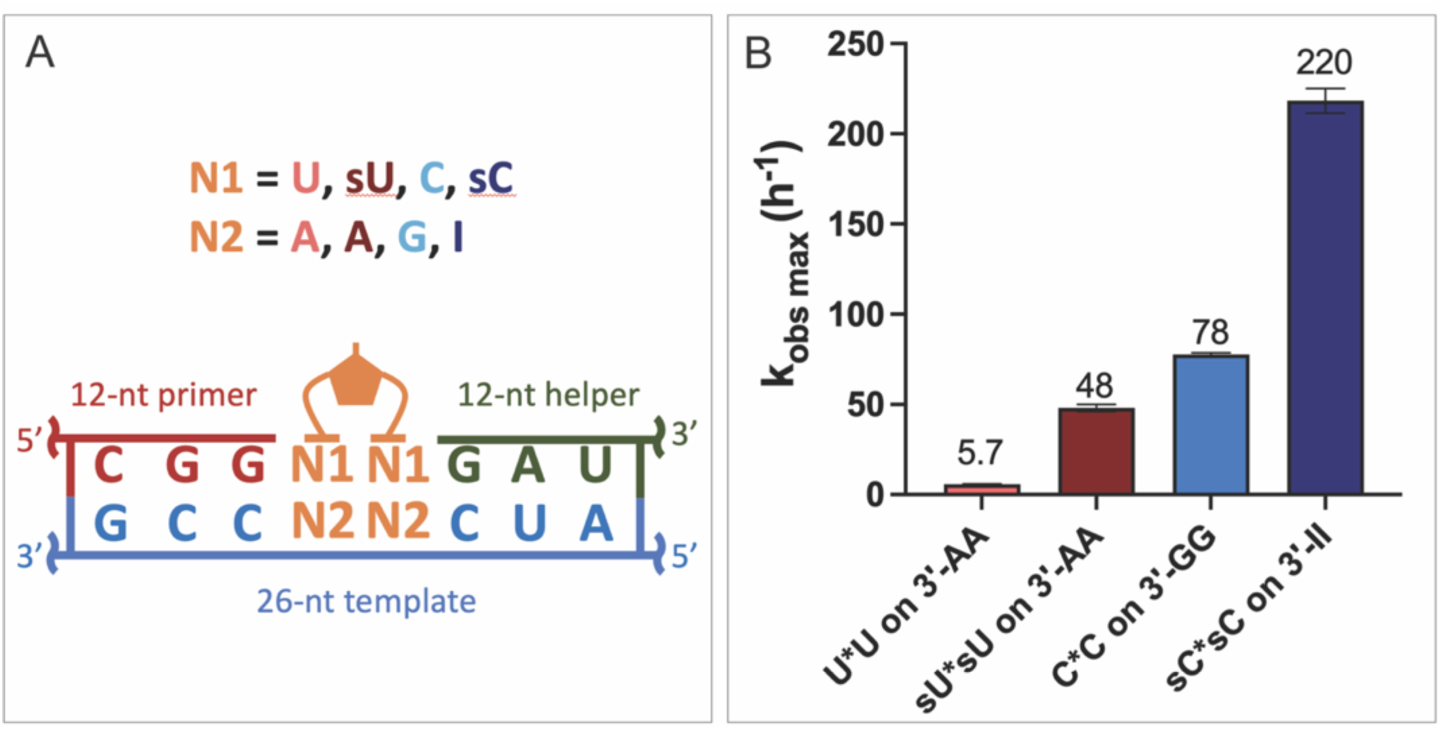
Primer extension with noncanonical bridged dinucleotides. (A) Scheme for the experiments. The template for sU*sU had two adenosines while the template for sC*sC had two inosines in the binding pocket. (B) Comparison of *k*_obs max_ between U*U, sU*sU, C*C, sC*sC in the sandwich model. For detailed description please see the SI.

## DISCUSSION

The recent discovery that the dominant pathway for nonenzymatic primer extension involves imidazolium-bridged dinucleotides as a covalent intermediate enabled us to reevaluate the kinetics of RNA template copying. Activated monomers first react with each other to form intermediate dimers, then bind to the template and react with the primer. Early studies of nonenzymatic RNA copying used activated mononucleotides, instead of purified intermediates; thus, the observed kinetics reflected the combination of distinct steps in the multi-step reaction. Nevertheless, these pioneering studies clearly showed that primer extension with C, G and A is much more efficient than with U.(8) Subsequent work showed that short activated helper oligonucleotides enable copying of mixed sequence templates, and facilitated the incorporation of A and U (19). Recent experiments using the purified dicytidine C*C intermediate provided for the first time a more accurate and comprehensive view of this multi-step chemical process.(5) Here, we have extended this analysis to a systematic study of the reaction kinetics of all ten canonical imidazolium-bridged dinucleotide intermediates. Our results show reasonably fast reaction rates (∼ 6 to 85 h^-1^) in all sixteen reactions when the intermediates are at saturating concentrations. However, the affinities of these intermediates for complementary templates exhibited a >7,000 fold range, from the very weak binding of U*U to the very tight binding of G*G. Given that equimolar activated monomers equilibrate to give roughly equimolar concentrations of bridged intermediates, it is clear that previous efforts to copy mixed sequence templates failed due to the extremely sub-saturating concentrations of the most weakly binding intermediates (U*U, U*A, A*C and U*G). Additional experiments will be required to see if adjusting the concentrations of monomers leads to enhanced copying of mixed sequence templates.

Our study only compared the reaction rates of each intermediate when they are sandwiched between guanosines at the end of the primer and the beginning of the downstream helper oligonucleotide. When stacking against other nucleobases, the reaction rates and affinities of the intermediates are likely to differ. In particular, the weak stacking of intermediates with the 3′-terminal U of a primer, or of bridged intermediates with a U in the downstream position, could make it difficult to reach saturating concentrations of intermediate at specific template positions, and could thus contribute to the difficulty of copying arbitrary template sequences. Thus other ways of modulating the affinity of bridged intermediates for the template may be needed to enable efficient sequence-general copying.

Our studies show that the efficiency of nonenzymatic primer extension with imidazolium-bridged dinucleotide intermediates is determined in part by the affinity of the dinucleotide for the primer/template/helper complex, but also by the sugar pucker of the incoming nucleotide. The affinities of the dinucleotide intermediates are affected by both the Watson-Crick base pairing and by the stacking interactions with neighboring nucleotides. Interestingly, the highest affinity intermediates also show the fastest reaction rates, even at saturating concentrations. While the reasons for this phenomenon remain unknown, we suggest that a series of strong interactions may stabilize the optimal configuration of the bound dinucleotides for reaction with the primer. It is also possible that weak interactions with the template-adjacent nucleotide (e.g. with U) may allow that nucleotide to transiently become unpaired to the template, even though the dinucleotide as a whole remains bound. The effect of sugar pucker in the imidazolium-bridged dinucleotide was unexpected, but emerged from the finding that the superior reactivity of imidazolium-bridged dinucleotides in which N1 was C correlated with the stronger preference of C for the C3ʹ-endo configuration, both as a monomer and in the intermediate. The preference of C for the C3ʹ-endo configuration has been previously reported.(20, 24) By measuring the rate of primer extension with G*G analogs with different sugar 2ʹ-substituents, we confirmed that the C3ʹ-conformation of the dinucleotide intermediate contributes to a faster rate of primer extension. The reason for this effect is not immediately obvious, since the sugars of the N1 and N2 nucleotides are remote from the site of the reaction. However, we suggest that the conformation of the sugar of the N1 nucleotide may influence the position of the reactive phosphate, relative the primer 3ʹ-hydroxyl.

The wide range of observed reaction rates and affinities among the ten different imidazolium-bridged dinucleotide intermediates clearly contributes to the difficulty of copying mixed sequence RNA templates. We asked whether this problem could potentially be overcome by employing either (i) noncanonical nucleotides, or (ii) activated short helper oligonucleotides. These potential solutions have been proposed before,(19, 25) but without recognition of the role of bridged-intermediates. Thiolated pyrimidine nucleotides have a strong preference for the C3ʹ-endo sugar conformation, and our experiments show that the sU*sU and sC*sC bridged dinucleotides exhibit significantly faster reaction rates than the canonical U*U and C*C intermediates. This is of particular interest given that both sC and sU can be generated form the prebiotic Sutherland pathway to the pyrimidine ribonucleotides.(26) As an alternative approach, we also prepared a series of monomer-bridged-oligonucleotides and compared their primer extension reaction rates with that of a standard bridged dinucleotide. Monomer bridged to a trimer exhibited a template affinity three orders of magnitude stronger than that of a simple bridged dinucleotide. In a prebiotic RNA replication scenario in which many short oligonucleotides were present (as in the virtual circular genome model)(27), and could become activated, such monomer-bridged oligonucleotide species would form naturally, and may have played a key role in enabling nonenzymatic RNA replication. We are currently examining the efficiency and fidelity of nonenzymatic primer extension in the presence of both noncanonical nucleotides and monomer-bridged-oligonucleotide species. We hope that these studies will lead to prebiotically realistic scenarios under which efficient nonenzymatic replication may take place.

## SUPPLEMENTARY DATA

Supplementary data are available at NAR online.

## ACKNOWLEDGEMENTS

We thank Dr. Marco Todisco, Dr. Li Li, Dr. Daniel Duzdevich, Dr. Seohyun Chris Kim and Lydia Pazienza for helpful discussion and technical assistance. We also thank all members of the Szostak laboratory for helpful feedback.

