## Supplementary Information for "Kinetic Explanations for the Sequence Biases Observed in the Nonenzymatic Copying of RNA Templates"

**1 Materials and Methods**
- **1.1 Abbreviations**
- **1.2 General information**
- **1.3 Procedure for homo-dinucleotide intermediate synthesis**
  - 1.3.1 1,3-di-(cytidine-5′phosphoryl)-2-aminoimidazolium (**C*C**)
  - 1.3.2 1,3-di-(guanosine-5′phosphoryl)-2-aminoimidazolium (**G*G**)
  - 1.3.3 1,3-di-(uridine-5′phosphoryl)-2-aminoimidazolium (**U*U**)
  - 1.3.4 1,3-di-(adenosine-5′phosphoryl)-2-aminoimidazolium (**A*A**)
  - 1.3.5 1,3-di-(2′-deoxyguanosine-5′phosphoryl)-2-aminoimidazolium (**DNA G*G**)
  - 1.3.6 1,3-di-(2′-deoxy-2′-fluoroguanosine-5′phosphoryl)-2-aminoimidazolium (**FRNA G*G**)
  - 1.3.7 1,3-di-(2′-O-Methylguanosine-5′phosphoryl)-2-aminoimidazolium (**2′-OMe G*G**)
  - 1.3.8 1,3-di-(2'-O, 4'-C methylene-guanosine-5′phosphoryl)-2-aminoimidazolium (**LNA G*G**)
  - 1.3.9 1,3-di-(2-thiouridine-5′phosphoryl)-2-aminoimidazolium (**sU*sU**)
  - 1.3.10 1,3-di-(2-thiocytidine-5′phosphoryl)-2-aminoimidazolium (**sC*sC**)
- **1.4 Procedure for hetero-dinucleotide intermediate synthesis**
  - 1.4.1 1-(cytidine-5′-phosphoryl)-3-(uridine-5′-phosphoryl)-2-aminoimidazolium (**C*U**)
  - 1.4.2 1-(cytidine-5′-phosphoryl)-3-(adenosine-5′-phosphoryl)-2-aminoimidazolium (**C*A**)
  - 1.4.3 1-(adenosine-5′-phosphoryl)-3-(uridine-5′-phosphoryl)-2-aminoimidazolium (**A*U**)
  - 1.4.4 1-(guanosine -5′-phosphoryl)-3-(uridine-5′-phosphoryl)-2-aminoimidazolium **(G*U**)
  - 1.4.5 1-(guanosine-5′-phosphoryl)-3-(adenosine-5′-phosphoryl)-2-aminoimidazolium (**G*A**)
  - 1.4.6 1-(guanosine-5′-phosphoryl)-3-(cytidine-5′-phosphoryl)-2-aminoimidazolium (**G*C**)
- **1.5 Oligonucleotide synthesis**
  - 1.5.1 Primers, templates, helpers, and complementary DNAs
  - 1.5.2 Monomer-bridged-oligonucleotide intermediates
- **1.6 Measurement of hetero-dinucleotide hydrolysis rate for Figure S1**
- **1.7 Primer extension reaction**
  - 1.7.1 Measuring the kinetics parameters in sandwich system for Figures 2, 3, 4, 8, S2, S3 and S6
  - 1.7.2 Measuring the kinetics parameters of modified guanosine imidazolium bridged dinucleotides for Figure 6, S3
  - 1.7.3 Measuring the kinetics parameters of monomer-bridged-oligo for Figure 7, S5

**2 Supplementary Figures and Tables**

### 1 Materials and Methods

#### 1.1 Abbreviation

2-AI, 2-aminoimidazole
CV, column volume
DIPEA, *N,N*-diisopropylethylamine
DMSO, dimethyl sulfoxide
DPDS, 2,2′-dipyridyldisulfide
EDTA, ethylenediaminetetraacetic acid
HOAt, 1-Hydroxy-7-azabenzotriazole
HPLC, high-performance liquid chromatography
HRMS, high-resolution mass spectrometry
LC-MS, liquid chromatography-mass spectrometry
NMR, nuclear magnetic resonance
PAGE, polyacrylamide gel electrophoresis
Q-TOF, quadrupole time-of-flight
TEA, triethylamine
TEAB, triethylammonium bicarbonate
TPP, triphenylphosphine
Tris, tris(hydroxymethyl)aminomethane
*, 2-aminoimidazole bridge

#### 1.2 General information

All chemicals were purchased from Sigma-Aldrich (St. Louis, MO) and used without purification unless otherwise noted. All ribonucleoside 5′-monophosphate free-acid compounds were purchased from MP Biotechnology (Solon, OH). 2,2′-Dipropyl disulfide was purchased from Combi-Blocks (San Diego, CA). 1-Hydroxy-7-azabenzotriazole was purchased from Creosalus (Louisville, KY). 2-thiouridine was purchased from Cayman Chemical (Ann Arbor, MI). 2-thiocytidine was purchased from Berry&Associates (Dexter, MI). Deuterated solvents were purchased from Cambridge Isotope Laboratories (Tewksbury, MA). Dmf-G-LA-CE Phosphoramidite was purchased from Glen Research (Sterling, VA). Reverse phase flash chromatography was performed using a prepacked RediSep Rf Gold C18Aq 50 g column from Teledyne Isco (Lincoln, NE). Preparatory-scale high performance liquid chromatography (HPLC) was carried out on an Agilent 1290 HPLC system, equipped with a preparative-scale Agilent ZORBAX Eclipse-XDB C18 column (21.2x250mm, 7 µm particle size) for reversed-phase chromatography.

$^{1}$H, and $^{31}$P spectra were acquired on a Varian Oxford AS-400 NMR spectrometer (400 MHz for $^{1}$H, 162 MHz for $^{31}$P). Chemical shifts are reported in parts per million (ppm) values on the δ scale. $^{1}$H NMR was referenced using DSS as internal standard (0 ppm at 25 ◦C). All NMR spectra were recorded at 25 °C. Data are reported as follows: chemical shift, multiplicity (s = singlet, d = doublet, t = triplet, q = quadruplet, m = multiplet), integration, and coupling constants. HRMS was carried out on an Agilent 1200 HPLC coupled to an Agilent 6230 TOF mass spectrometer.

#### 1.3 Procedure for homo-dinucleotide intermediate synthesis

##### 1.3.1 1,3-di-(cytidine-5′phosphoryl)-2-aminoimidazolium (C*C)

The homo-dinucleotide **C*C** was synthesized according to the literature procedure. (1, 2)
**$^{1}$H NMR** (400 MHz, $D_2O$) δ 7.64 (d, *J* = 7.6 Hz, 2H), 6.93 (t, *J* = 2.0 Hz, 2H), 6.02 (d, *J* = 7.6 Hz, 2H), 5.82 (d, *J* = 3.2 Hz, 2H), 4.25 – 4.11 (m, 10H). Peaks corresponding to residual TEAB observed at 3.20 and 1.27 ppm. **$^{31}$P NMR** (162 MHz, $D_2O$) δ -12.85. **HRMS** (Q-TOF) m/z: $[M - H]^-$ Calcd. for $C_{21}H_{28}N_9O_{14}P_2$ 692.1231; Found: 692.1295.

##### 1.3.2 1,3-di-(guanosine-5′phosphoryl)-2-aminoimidazolium (G*G)

The homo-dinucleotide **G*G** was synthesized according to the literature procedure. (3)
**$^{1}$H NMR** (400 MHz, $D_2O$) δ 7.83 (s, 2H), 6.70 – 6.68 (m, 2H), 5.75 (d, *J* = 5.2 Hz, 2H), 4.65 (t, *J* = 5.3 Hz, 2H), 4.39 (t, *J* = 5.0 Hz, 2H), 4.14 – 4.07 (m, 4H), 4.03 – 3.96 (m, 2H). Peaks corresponding to residual TEAB observed at 3.19 and 1.27 ppm. **$^{31}$P NMR** (162 MHz, $D_2O$) δ -12.85. **HRMS** (Q-TOF) m/z: $[M - H]^-$ Calcd. for $C_{23}H_{28}N_{13}O_{14}P_2$ 772.1354; Found: 772.1411.

##### 1.3.3 1,3-di-(uridine-5′phosphoryl)-2-aminoimidazolium (U*U)

The homo-dinucleotide **U*U** was synthesized using the same procedure as **C*C** in 1.3.1.
**$^{1}$H NMR** (400 MHz, $D_2O$) δ 7.64 (d, *J* = 8.1 Hz, 2H), 6.95 (t, *J* = 2.0 Hz, 2H), 5.88 (d, *J* = 8.1 Hz, 2H), 5.82 (d, *J* = 4.3 Hz, 2H), 4.27 (t, *J* = 4.9 Hz, 2H), 4.21 (t, *J* = 5.3 Hz, 2H), 4.19 – 4.16 (m, 4H), 4.15 – 4.11 (m, 2H). Peaks corresponding to residual TEAB observed at 3.19 and 1.27 ppm. **$^{31}$P NMR** (162 MHz, $D_2O$) δ -12.90. **HRMS** (Q-TOF) m/z: $[M - H]^-$ Calcd. for $C_{21}H_{26}N_7O_{16}P_2$ 694.0911; Found: 694.0932.

##### 1.3.4 1,3-di-(adenosine-5′phosphoryl)-2-aminoimidazolium (A*A)

The homo-dinucleotide **A*A** was synthesized using the same procedure as **C*C** in 1.3.1.
**$^{1}$H NMR** (400 MHz, $D_2O$) δ 8.12 (s, 2H), 8.07 (s, 2H), 6.74 – 6.71 (m, 2H), 5.87 (d, *J* = 4.8 Hz, 2H), 4.58 (t, *J* = 5.0 Hz, 2H), 4.37 (t, *J* = 5.0 Hz, 2H), 4.16 – 4.06 (m, 4H), 4.05 – 3.98 (m, 2H). Peaks corresponding to residual TEAB observed at 3.20 and 1.27 ppm. **$^{31}$P NMR** (162 MHz, $D_2O$) δ -12.81. **HRMS** (Q-TOF) m/z: $[M - H]^-$ Calcd. for $C_{23}H_{28}N_{13}O_{12}P_2$ 740.1456; Found: 740.1553.

##### 1.3.5 1,3-di-(2′-deoxyguanosine-5′phosphoryl)-2-aminoimidazolium (DNA G*G)

The homo-dinucleotide **DNA G*G** was synthesized from the 2′-deoxyguanosine 5′-monophosphate using the same procedure as **G*G** in 1.3.2.
**$^{1}$H NMR** (400 MHz, $D_2O$) δ 7.84 (s, 2H), 6.63 – 6.60 (m, 2H), 6.13f (t, *J* = 6.6 Hz, 2H), 4.62 – 4.56 (m, 2H), 4.10 – 4.02 (m, 4H), 3.97 (dt, *J* = 13.4, 6.6 Hz, 2H), 2.72 (dt, *J* = 13.4, 6.6 Hz, 2H), 2.46 (ddd, *J* = 14.0, 6.7, 4.3 Hz, 2H). Peaks corresponding to residual TEAB observed at 3.19 and 1.29 ppm. **$^{31}$P NMR** (162 MHz, $D_2O$) δ -12.82. **HRMS** (Q-TOF) m/z: $[M - H]^-$ Calcd. for $C_{23}H_{28}N_{13}O_{12}P_2$ 740.1461; Found: 740.1523.

##### 1.3.6 1,3-di-(2′-deoxy-2′-fluoroguanosine-5′phosphoryl)-2-aminoimidazolium (FRNA G*G)

The 2′-deoxy-2′-fluoroguanosine 5′-monophosphate was synthesized from the corresponding nucleosides by the procedure of Yoshikawa.(4) The homo-dinucleotide **FRNA G*G** was

synthesized from the 2′-deoxy-2′-fluoroguanosine 5′-monophosphate using the same procedure as **G*G** in 1.3.2.
**$^{1}$H NMR** (400 MHz, $D_2O$) δ 7.74 (s, 2H), 6.56 (s, 2H), 5.09 (dd, *J* = 19.5, 1.6 Hz, 2H), 5.43 – 5.23 (m, 2H), 4.69 (ddd, *J* = 21.6, 8.1, 4.7 Hz, 2H), 4.14 (dt, *J* = 12.0, 4.7 Hz, 2H), 4.10 – 3.95 (m, 4H). Peaks corresponding to residual TEAB observed at 3.19 and 1.27 ppm. **$^{31}$P NMR** (162 MHz, $D_2O$) δ -10.25. **HRMS** (Q-TOF) m/z: $[M - H]^-$ Calcd. for $C_{23}H_{26}F_2N_{13}O_{12}P_2$ 776.1273; Found: 740.1335.

##### 1.3.7 1,3-di-(2′-O-Methylguanosine-5′phosphoryl)-2-aminoimidazolium (2′-OMe G*G)

The 2′-OMe-guanosine 5′-monophosphate was synthesized from the corresponding nucleosides by the procedure of Yoshikawa.(4) The homo-dinucleotide **2′-OMe G*G** was synthesized from the 2′-O-methylguanosine monophosphate using the same procedure as **G*G** in 1.3.2.
**$^{1}$H NMR** (400 MHz, $D_2O$) δ 7.84 (s, 2H), 6.66 (s, 2H), 5.80 (d, *J* = 5.0 Hz, 2H), 4.53 (t, *J* = 4.9 Hz, 2H), 4.34 (t, *J* = 5.1 Hz, 2H), 4.08 (dt, *J* = 11.2, 4.3 Hz, 2H), 4.01 – 3.97 (m, 2H), 3.94 – 3.86 (m, 2lH), 3.43 (s, 6H). Peaks corresponding to residual TEAB observed at 3.19 and 1.27 ppm. **$^{31}$P NMR** (162 MHz, $D_2O$) δ -12.73. **HRMS** (Q-TOF) m/z: $[M - H]^-$ Calcd. for $C_{25}H_{32}N_{13}O_{14}P_2$ 800.1672; Found: 800.1723.

##### 1.3.8 1,3-di-(2'-O, 4'-C methylene-guanosine-5′phosphoryl)-2-aminoimidazolium (LNA G*G)

Since the locked nucleic acid (LNA) guanosine nucleoside is not commercially available, we resorted to purchasing the dmf-G-LA-CE Phosphoramidite (Glen Research) which we have coupled to the Universal Support III PS (Glen Research) using a MerMade 6 DNA/RNA synthesizer (Bioautomation, Plano, TX). After cleavage from the resin and base deprotection with concentrated ammonia solution and purification by reverse phase flash chromatography, the nucleoside was phosphorylated using the Yoshikawa protocol.(4) The homo-dinucleotide **LNA G*G** was synthesized from the locked nucleic acid guanosine monophosphate using the same procedure as **G*G** in 1.3.2.
$^{1}$H NMR (400 MHz, $D_2O$) δ 7.65 (s, 2H), 6.90 (t, *J* = 1.9 Hz, 2H), 5.61 (s, 2H), 4.52 (s, 2H), 4.41 (s, 2H), 4.39 – 4.23 (m, 4H), 4.03 (d, *J* = 8.6 Hz, 2H), 3.84 (d, *J* = 8.6 Hz, 2H). Peaks corresponding to residual TEAB observed at 3.20 and 1.27 ppm. **$^{31}$P NMR** (162 MHz, $D_2O$) δ -12.84. **HRMS** (Q-TOF) m/z: $[M - H]^-$ Calcd. for $C_{25}H_{28}N_{13}O_{14}P_2$ 796.1359; Found: 796.1422.

##### 1.3.9 1,3-di-(2-thiouridine-5′phosphoryl)-2-aminoimidazolium (sU*sU)

The synthesis of the homo-dinucleotide **sU*sU** began by first synthesizing the 2-thiouridine 5′-monophosphate (**sUMP**). To a pre-chilled solution of trimethylphosphate (9 mL) were added 2-thiouridine (270 mg), $POCl_3$ (0.35 mL), and $H_2O$ (7.5 uL). The resultant solution was stirred at 0°C until complete solubilization. Then four separate portions of DIPEA (94 uL each) were added dropwise into the stirred solution with 20 min intervals at 0°C. The reaction mixture was allowed to stir for an additional 20 min until quenched by 1M TEAB (20 mL, pH 7.6). The product was partially purified by reverse phase flash chromatography using aqueous 25 mM TEAB (pH 7.5) over 5 CVs with a flow rate of 40 mL/min. Fractions containing product were pooled and lyophilized. The resultant syrup was dissolved in water and repurified by reverse phase flash chromatography using gradient elution between (A) aqueous 25 mM TEAB (pH 7.5) and (B) acetonitrile. The sample was eluted between 0% and 10% B over 10 CVs with a flow rate of 40 mL/min. Fractions containing **sUMP** were pooled and lyophilized.

Part of the **sUMP** was activated to form the activated monomer 2-thiouridine 5′-phosphoryl-(2-aminoimidazole) (**2-AI-sU**). **sUMP** (385 umol), 2-aminoimidazole hydrochloride (460 mg), and TPP (1010 mg) were dissolved in DMSO (10 mL) and TEA (2 mL). The reaction mixture was added DPDS (850 mg), and it was stirred at room temperature for 2 hours. After the reaction was completed, the mixture was precipitated by adding acetone (80 ml) and saturated $NaClO_4$ in acetone (4 mL). The precipitate was pelleted by centrifugation (3000 rpm, 10 min) and washed with 1:1 mixture of acetone:diethylether. After decanting the solvent, the pellet was dried under vacuum, resuspended in deionized water, and purified by reverse phase flash chromatography using gradient elution between (A) aqueous 2 mM TEAB buffer (pH 8.0) and (B) acetonitrile. The sample was eluted between 0% and 10% B over 10 CVs with a flow rate of 40 mL/min. Fractions containing **2-AI-sU** were pooled and lyophilized at -20 ◦C.
**$^1$H NMR** (400 MHz, $D_2O$) δ 7.86 (d, *J* = 7.9 Hz, 1H), 6.79 (t, *J* = 1.7 Hz, 1H), 6.73 (d, *J* = 3.3 Hz, 1H), 6.62 (t, *J* = 2.0 Hz, 1H), 6.08 (d, *J* = 7.9 Hz, 1H), 4.31 (dd, *J* = 5.0, 3.4 Hz, 1H), 4.26 – 4.20 (m, 2H), 4.18 – 4.14 (m, 1H), 4.13 – 4.07 (m, 1H). Peaks corresponding to residual TEAB observed at 3.19 and 1.27 ppm. **$^{31}$P NMR** (162 MHz, $D_2O$) δ -10.97. **HRMS** (Q-TOF) m/z: $[M - H]^-$ Calcd. for $C_{12}H_{15}N_5O_7PS$ 404.0435; Found: 404.0589.
Next, the **2-AI-sU** was used to activate **sUMP** to form the desired homo-dinucleotide **sU*sU**. **2-AI-sU** (64 umol), **sUMP** (74 umol), and TPP (503 mg) were dissolved in DMSO (5 mL) and TEA (1 mL). The reaction mixture was added DPDS (423 mg), and it was stirred at room temperature for 15 min. The mixture was then precipitated by adding acetone (40 ml) and saturated $NaClO_4$ in acetone (2 mL). The precipitate was pelleted by centrifugation (3000 rpm, 10 min) and washed with 1:1 mixture of acetone:diethylether. After decanting the solvent, the pellet was dried under vacuum, resuspended in deionized water, and purified by reverse phase flash chromatography using gradient elution between (A) aqueous 2 mM TEAB buffer (pH 7.5) and (B) acetonitrile. The sample was eluted between 0% and 10% B over 25 CVs with a flow rate of 40 mL/min. Fractions containing **sU*sU** were adjusted to pH 8.0 and lyophilized at -20 ◦C.
**$^1$H NMR** (400 MHz, $D_2O$) δ 7.75 (d, *J* = 8.1 Hz, 2H), 7.03 (t, *J* = 2.1 Hz, 2H), 6.45 (d, *J* = 2.3 Hz, 2H), 6.10 (d, *J* = 8.1 Hz, 2H), 4.33 – 4.21 (m, 6H), 4.20 – 4.11 (m, 4H). Peaks corresponding to residual TEAB observed at 3.19 and 1.27 ppm. **$^{31}$P NMR** (162 MHz, $D_2O$) δ -12.94. **HRMS** (Q-TOF) m/z: $[M - H]^-$ Calcd. for $C_{21}H_{26}N_7O_{14}P_2S_2$ 726.0460; Found: 726.0669.

##### 1.3.10 1,3-di-(2-thiocytidine-5′phosphoryl)-2-aminoimidazolium (sC*sC)

The activated monomer **2-AI-sC** was synthesized from 2-thiocytidine first using the same procedure as **2-AI-sU** in 1.3.9.
**$^1$H NMR** (400 MHz, $D_2O$) δ 7.98 (d, *J* = 7.7 Hz, 1H), 6.80 (t, *J* = 1.5 Hz, 1H), 6.72 (s, 1H), 6.66 (d, *J* = 2.4 Hz, 1H), 6.64 (t, *J* = 1.8 Hz, 1H), 4.32 (dd, *J* = 4.9, 2.4 Hz, 1H), 4.29 – 4.21 (m, 2H), 4.19 – 4.07 (m, 2H). Peaks corresponding to residual TEAB observed at 3.19 and 1.27 ppm. **$^{31}$P NMR** (162 MHz, $D_2O$) δ -10.71. **HRMS** (Q-TOF) m/z: $[M - H]^-$ Calcd. for $C_{12}H_{16}N_6O_6PS$ 403.0595; Found: 403.0778.
Then the homo-dinucleotide **sC*sC** was synthesized from **2-AI-sC** using the same procedure as **sU*sU**.
**$^1$H NMR** (400 MHz, $D_2O$) δ 7.82 (d, *J* = 7.4 Hz, 2H), 7.01 (s, 2H), 6.52 (s, 2H), 6.29 (d, *J* = 7.6 Hz, 2H), 4.42 – 4.00 (m, 10H). Peaks corresponding to residual TEAB observed at 3.20 and 1.27 ppm. **$^{31}$P NMR** (162 MHz, $D_2O$) δ -12.89. **HRMS** (Q-TOF) m/z: $[M - H]^-$ Calcd. for $C_{21}H_{28}N_9O_{12}P_2S_2$ 724.0780; Found: 724.0971.

#### 1.4 Procedure for hetero-dinucleotide intermediate synthesis

##### 1.4.1 1-(cytidine-5′-phosphoryl)-3-(uridine-5′-phosphoryl) -2-aminoimidazolium (C*U)

The synthesis of the hetero-dinucleotide **C*U** began by first synthesizing two monomers, cytidine 5′-phosphoryl-(1-hydroxy-7-azabenzotriazole) (**C-OAt**) and uridine 5′-phosphoryl-(2-aminoimidazole) (**2-AI-U**).

Cytidine 5′-monophosphate (100 mg), HOAt (220 mg), and TPP (850 mg) were dissolved in DMSO (4 mL) and TEA (1 mL). DPDS (700 mg) was added to the reaction mixture, which was stirred at room temperature for 2 hours. After the reaction was completed, the mixture was precipitated by adding acetone (40 ml) and saturated $NaClO_4$ in acetone (2 mL). The precipitate was pelleted by centrifugation (3000 rpm, 10 min) and washed with 1:1 mixture of acetone:diethylether. After decanting the solvent, the pellet was dried under vacuum, resuspended in deionized water and purified by reverse phase flash chromatography using gradient elution between (A) aqueous 2 mM TEAB buffer (pH 8.0) and (B) acetonitrile. The sample was eluted between 0% and 10% B over 10 CVs with a flow rate of 40 mL/min. Fractions containing **C-OAt** were pooled and lyophilized at -20 ◦C.

Uridine 5′-monophosphate (80 mg), 2-aminoimidazole hydrochloride (160 mg), and TPP (520 mg) were dissolved in DMSO (5 mL) and TEA (1 mL). The reaction mixture was added DPDS (400 mg), and it was stirred at room temperature for 2 hours. After the reaction was completed, the mixture was precipitated by adding acetone (40 ml) and saturated $NaClO_4$ in acetone (2 mL). The precipitate was pelleted by centrifugation (3000 rpm, 10 min) and washed with 1:1 mixture of acetone:diethylether. After decanting the solvent, the pellet was dried under vacuum, resuspended in deionized water and purified by reverse phase flash chromatography using gradient elution between (A) aqueous 2 mM TEAB buffer (pH 8.0) and (B) acetonitrile. The sample was eluted between 0% and 10% B over 10 CVs with a flow rate of 40 mL/min. Fractions containing **2-AI-U** were pooled and lyophilized at -20 ◦C.

The lyophilized **C-OAt** and **2-AI-U** were dissolved in deionized water (5 mL) and adjusted pH to 8.0 by NaOH/HCl, then stirred for 1 hour at room temperature. After the reaction completed, the mixture was purified by reverse phase HPLC over a C18 column with (A) 2 mM TEAB buffer (pH 8.0) and (B) acetonitrile. The sample was eluted between 2% and 8% B over 20 CVs with a flow rate of 15 mL/min. Fractions containing product were pooled and lyophilized at -20 ◦C.

**$^1$H NMR** (400 MHz, $D_2O$) δ 7.65 (d, *J* = 8.0 Hz, 1H), 7.64 (d, *J* = 7.5 Hz, 1H), 6.94 (s, 2H), 6.02 (d, *J* = 7.5 Hz, 1H), 5.88 (d, *J* = 8.1 Hz, 1H), 5.84 (d, *J* = 2.7 Hz, 1H), 5.80 (d, *J* = 4.2 Hz, 1H), 4.28 – 4.24 (m, 1H), 4.23 – 4.11 (m, 9H). Peaks corresponding to residual TEAB observed at 3.20 and 1.27 ppm. **$^{31}$P NMR** (162 MHz, $D_2O$) δ -12.88 (d, *J* = 13.6 Hz). **HRMS** (Q-TOF) m/z: Calcd. for $C_{22}H_{27}N_{10}O_{14}P_2$ $[M - H]^-$ 693.1071; Found: 693.1154

##### 1.4.2 1-(cytidine-5′-phosphoryl)-3-(adenosine-5′-phosphoryl)-2-aminoimidazolium (C*A)

The hetero-dinucleotide **C*A** was prepared following similar procedure as **C*U** in section 1.4.1.

**$^1$H NMR** (400 MHz, $D_2O$) δ 8.22 (s, 1H), 7.46 (d, *J* = 7.7 Hz, 1H), 6.80 (d, *J* = 9.6 Hz, 2H), 6.00 (d, *J* = 4.9 Hz, 1H), 5.89 (d, *J* = 7.5 Hz, 1H), 5.70 (d, *J* = 2.6 Hz, 1H), 4.67 (t, *J* = 5.0 Hz, 1H), 4.44 (t, *J* = 4.9 Hz, 1H), 4.24 – 4.06 (m, 7H), 3.99 – 3.95 (m, 1H). Peaks corresponding to residual TEAB observed at 3.19 and 1.27 ppm. **$^{31}$P NMR** (162 MHz, $D_2O$) δ -12.83 (d, *J* = 15.5 Hz). **HRMS** (Q-TOF) m/z: $[M - H]^-$ Calcd. for $C_{23}H_{28}N_{13}O_{12}P_2$ 716.1343; Found: 716.1418.

**1.4.3 1-(adenosine-5′-phosphoryl)-3-(uridine-5′-phosphoryl)-2-aminoimidazolium(A*U)**

The hetero-dinucleotide **A*U** was prepared following similar procedure as **C*U** in section 1.4.1. $^{1}$**H NMR** (400 MHz, $D_2O$) δ 8.23 (s, 1H), 8.22 (s, 1H), 7.49 (d, *J* = 8.1 Hz, 1H), 6.80 (d, *J* = 15.5 Hz, 2H), 6.00 (d, *J* = 4.9 Hz, 1H), 5.80 (d, *J* = 8.1 Hz, 1H), 5.65 (d, *J* = 4.1 Hz, 1H), 4.67 (t, *J* = 5.2 Hz, 1H), 4.45 (t, *J* = 4.9 Hz, 1H), 4.20 – 4.25 (m, 2H), 4.19 – 4.12 (m, 3H), 3.81 – 3.77 (m, 3H). Peaks corresponding to residual TEAB observed at 3.19 and 1.27 ppm. $^{31}$**P NMR** (162 MHz, $D_2O$) δ -12.88 (d, *J* = 34.9 Hz). **HRMS** (Q-TOF) m/z: Calcd. for $C_{22}H_{27}N_{10}O_{14}P_2$ $[M - H]^-$ 717.1184; Found: 717.1184.

**1.4.4 1-(guanosine -5′-phosphoryl)-3-(uridine-5′-phosphoryl)-2-aminoimidazolium (G*U)**

The synthesis of the hetero-dinucleotide **G*U** began by first synthesizing two monomers, guanosine 5′-phosphoryl-(1-hydroxy-7-azabenzotriazole) (**G-OAt**) and uridine 5′-phosphoryl-(2-aminoimidazole) (**2-AI-U**).

Guanosine 5′-monophosphate (100 mg), HOAt (200 mg), and 2-aminoimidazole hydrochloride (200 mg) were dissolved in water (8 mL). The solution was vortexed and sonicated until completely homogenized, then flash frozen in liquid nitrogen and lyophilized. Once dry, the solid was added to TPP (750 mg), and dissolved in DMSO (4 mL) and TEA (1 mL). To the reaction mixture was added DPDS (625 mg), with stirring at room temperature for 2 hours. After the reaction was complete, the mixture was precipitated by adding acetone (40 ml) and saturated $NaClO_4$ in acetone (2 mL). The precipitate was pelleted by centrifugation (3000 rpm, 10 min) and washed with 1:1 mixture of acetone:diethylether. After decanting the solvent, the pellet was dried under vacuum, resuspended in deionized water and purified by reverse phase flash chromatography using gradient elution between (A) aqueous 2 mM TEAB buffer (pH 8.0) and (B) acetonitrile. The sample was eluted between 0% and 10% B over 10 CVs with a flow rate of 40 mL/min. Fractions containing **G-OAt** were pooled and lyophilized at -20 ◦C.

Uridine 5′-monophosphate (80 mg), 2-aminoimidazole hydrochloride (160 mg), and TPP (520 mg) were dissolved in DMSO (4 mL) and TEA (1 mL). The reaction mixture was added DPDS (400 mg), and it was stirred at room temperature for 2 hours. After the reaction was completed, the mixture was precipitated by adding acetone (40 ml) and saturated $NaClO_4$ in acetone (2 mL). The precipitate was pelleted by centrifugation (3000 rpm, 10 min) and washed with 1:1 mixture of acetone:diethylether. After decanting the solvent, the pellet was dried under vacuum, resuspended in deionized water and purified by reverse phase flash chromatography using gradient elution between (A) aqueous 2 mM TEAB buffer (pH 8.0) and (B) acetonitrile. The sample was eluted between 0% and 10% B over 10 CVs with a flow rate of 40 mL/min. Fractions containing **2-AI-U** were adjusted to pH 10.0 and lyophilized at -20 ◦C.

The lyophilized **G-OAt** and **2-AI-U** were dissolved in deionized water (5 mL) and adjusted pH to 8.0 by NaOH/HCl, then stirred for 1 hour at room temperature. After the reaction completed, the mixture was purified by reverse phase HPLC over a C18 column with (A) 2 mM TEAB buffer (pH 8.0) and (B) acetonitrile. The sample was eluted between 2% and 5% B over 30 CVs with a flow rate of 15 mL/min. Fractions containing product were adjusted to pH 8.0 and lyophilized at -20 ◦C. $^{1}$**H NMR** (400 MHz, $D_2O$) δ 7.88 (s, 1H), 7.55 (d, *J* = 8.1 Hz, 1H), 6.81 (d, *J* = 13.7 Hz, 2H), 5.83 (d, *J* = 8.0 Hz, 1H), 5.80 (d, *J* = 5.4 Hz, 1H), 5.76 (d, *J* = 4.3 Hz, 1H), 4.67 (t, *J* = 5.2 Hz, 1H), 4.43 (t, *J* = 4.6 Hz, 1H), 4.24 – 4.16 (m, 5H), 4.05 – 4.00 (m, 3H). Peaks corresponding to residual TEAB observed at 3.19 and 1.27 ppm. $^{31}$**P NMR** (162 MHz, $D_2O$) δ -12.87 (d, *J* = 32.0 Hz). **HRMS** (Q-TOF) m/z: Calcd. for $C_{22}H_{27}N_{10}O_{15}P_2$ $[M - H]^-$ 733.1138; Found:733.1193.

##### 1.4.5 1-(guanosine-5′-phosphoryl)-3-(adenosine-5′-phosphoryl)-2-aminoimidazolium (G*A)

The hetero-dinucleotide **A*G** was prepared following similar procedure as **G*U** in section 1.4.4. The purification was performed using reverse phase HPLC over a C18 column with (A) 2 mM TEAB buffer (pH 8.0) and (B) acetonitrile. The sample was eluted between 2% and 8% B over 24 CVs with a flow rate of 15 mL/min. Fractions containing product were adjusted to pH 8.0 and lyophilized at -20 ˚C.

**$^1$H NMR** (400 MHz, $D_2O$) δ 8.19 (s, 1H), 8.14 (s, 1H), 7.77 (s, 1H), 6.69 (s, 2H), 5.95 (d, *J* = 4.9 Hz, 1H), 5.67 (d, *J* = 5.1 Hz, 1H), 4.64 (t, *J* = 5.1 Hz, 1H), 4.57 (t, *J* = 5.3 Hz, 1H), 4.41 (t, *J* = 4.9 Hz, 1H), 4.36 (t, *J* = 4.9 Hz, 1H), 4.17 – 4.04 (m, 3H), 4.03 – 3.92 (m, 3H). Peaks corresponding to residual TEAB observed at 3.19 and 1.27 ppm. **$^{31}$P NMR** (162 MHz, $D_2O$) δ -12.82. **HRMS** (Q-TOF) m/z: Calcd. for $C_{23}H_{28}N_{13}O_{13}P_2$ $[M - H]^-$ 756.1410; Found: 756.1535.

##### 1.4.6 1-(guanosine-5′-phosphoryl)-3-(cytidine-5′-phosphoryl)-2-aminoimidazolium (G*C)

The hetero-dinucleotide **C*G** was prepared following a similar procedure as **G*U** in section 1.4.4. The purification was performed using reverse phase HPLC over a C18 column with (A) 2 mM TEAB buffer (pH 8.0) and (B) acetonitrile. The sample was eluted between 2% and 6% B over 24 CVs with a flow rate of 15 mL/min. Fractions containing product were adjusted to pH 8.0 and lyophilized at -20 ◦C.

**$^1$H NMR** (400 MHz, $D_2O$) δ 7.89 (s, 1H), 7.58 (d, *J* = 7.6 Hz, 1H), 6.80 (q, *J* = 2.2 Hz, 2H), 6.00 (d, *J* = 7.5 Hz, 1H), 5.84 (d, *J* = 5.2 Hz, 1H), 5.83 (d, *J* = 3.3 Hz, 1H), 4.66 (t, *J* = 5.2 Hz, 1H), 4.44 (t, *J* = 4.9 Hz, 1H), 4.22 – 4.17 (m, 4H), 4.15 – 4.05 (m, 4H). Peaks corresponding to residual TEAB observed at 3.18 and 1.29 ppm. **$^{31}$P NMR** (162 MHz, $D_2O$) δ -12.82 (d, *J* = 14.6 Hz). **HRMS** (Q-TOF) m/z: Calcd. for $C_{22}H_{27}N_{10}O_{14}P_2$ $[M - H]^-$ 732.1293; Found: 732.1357.

#### 1.5 Oligonucleotide synthesis

##### 1.5.1 Primers, templates, helpers, and complementary DNAs

All oligonucleotides used in this study are listed in Table S1. Synthetic oligonucleotides were either purchased from Integrated DNA Technologies or prepared by solid phase synthesis using Expedite 8909 DNA/RNA synthesizer with phosphonamidites and reagents from ChemGenes (Wilmington, MA) and Glen Research (Sterling, MA). Oligonucleotides prepared in-house were deprotected and purified by Glen-Pak™ RNA purification cartridge (Sterling, MA).

##### 1.5.2 Monomer-bridged-oligo intermediates

The 5′-phosphorylated dimer pCG, trimer pCGC, and tetramer pCGCA oligonucleotides used in this study were prepared by solid-phase synthesis using MerMade 6 DNA/RNA synthesizer (Bioautomation, Plano, TX). The 5′-phosphorylated oligo nucleotides were deprotected and purified by reverse phase flash chromatography using reverse phase flash chromatography using gradient elution between (A) aqueous 25 mM TEAB buffer (pH 7.5) and (B) acetonitrile. The sample was eluted between 0% and 10% B over 10 CVs with a flow rate of 40 mL/min. Fractions containing the oligonucleotides were pooled and lyophilized at room temperature.

The oligonucleotide (26.2 umol), 2-aminoimidazole hydrochloride (125 mg), and TPP (275 mg) were dissolved in DMSO (4 mL) and TEA (1 mL). The reaction mixture was added DPDS (231 mg), and it was stirred at room temperature for 6 hours. After the reaction was completed, the mixture was precipitated by adding acetone (40 ml) and saturated $NaClO_4$ in acetone (2 mL). The precipitate was pelleted by centrifugation (3000 rpm, 10 min) and washed with 1:1 mixture of acetone:diethylether. After decanting the solvent, the pellet was dried under vacuum, resuspended in deionized water and purified by reverse phase flash chromatography using gradient elution between (A) aqueous 25 mM TEAB buffer (pH 7.5) and (B) acetonitrile. The sample was eluted between 0% and 10% B over 10 CVs with a flow rate of 40 mL/min. Fractions containing the activated oligonucleotide were pooled and lyophilized at -20 ◦C. **A-OAt** was synthesized following a similar procedure as **C-OAt** in 1.4.1. The activated oligonucleotide and 1.5 equiv. of the **A-OAt** were dissolved in deionized water (5 mL) and adjusted pH to 8.0 by NaOH/HCl, then stirred for 1 hour at room temperature. After the reaction completed, the mixture was purified by reverse phase HPLC over a C18 column with (A) 25 mM TEAB buffer (pH 7.5) and (B) acetonitrile. The sample was eluted between 0% and 10% B over 27 CVs with a flow rate of 40 mL/min. Fractions containing monomer-bridged-oligos were pooled and lyophilized at -20 ◦C.

#### 1.6 Measurement of hetero-dinucleotide degradation rate for Figure S1

Degradation of hetero-dinucleotides **A*C** and **A*U** were performed at 25°C in primer extension buffers either with magnesium anion (200 mM d11-Tris pH 8.0, 100 mM $MgCl_2$) or without magnesium anion (200 mM d11-Tris pH 8.0). The degradation experiments were initiated by adding 5 mM of the hetero-dinucleotide and monitored at 25°C by $^1H$ spectroscopy on a Varian 400 MHz NMR spectrometer (Oxford AS-400). The corresponding NMR peaks for each species were identified by the authentic NMR patterns in section 1.3 and 1.4. Peaks corresponding to residual TEAB at 3.19 and 1.27 ppm were used as the internal standard for integration and concentration calculations. The half-life was estimated based on the time point where the concentration of the hetero-dinucleotide **A*C** or **A*U** had dropped to approximately 2.5 mM.

#### 1.7 Primer extension reaction

##### 1.7.1 Measuring kinetic parameters in the sandwich system for Figures 2, 3, 4, 8, S2, S3 and S6

The primer-template-helper complexes were prepared in a solution containing 7.5 μM primer, 12.5 μM template, 17.5 μM helper, 50 mM Tris (pH 8.0), 50 mM NaCl and 1 mM EDTA (pH 8.0) by heating at 85°C for 30 s and slowly cooling down to 25°C at a rate of 0.1°C/s. The annealed product was diluted five-fold in primer extension reaction. Stock solutions of bridged dinucleotides/monomers used in the reaction were prepared freshly and adjusted to pH 8.0 immediately before the reaction.

At each time point, 0.5 μl of reaction sample was added to 25 μl quenching buffer, containing 25 mM EDTA, 1X TBE and 4 μM complementary DNA of template in formamide. Primer extension products were resolved by 20% (19:1) denaturing PAGE. The gel was scanned using a Typhoon 9410 scanner, and the bands were quantified using the ImageQuant TL software. The primers, templates, helpers, and complementary DNAs used in the primer extension assays are listed below:

| Bridged dinucleotide /monomer added | Primer | Template | Helper | Complimentary DNA |
|---|---|---|---|---|
| U*U | DL-30 | LA-121 | LA-111 | dCLA-121 |
| U*A | DL-30 | LA-115 | LA-111 | dCLA-115 |
| U*C | DL-30 | LA-120 | LA-111 | dCLA-120 |
| U*G | DL-30 | LA-109 | LA-111 | dCLA-109 |
| A*U | DL-30 | LA-116 | LA-111 | dCLA-116 |
| A*A | DL-30 | LA-122 | LA-111 | dCLA-122 |
| A*C | DL-30 | LA-117 | LA-111 | dCLA-117 |
| A*G | DL-30 | LA-113 | LA-111 | dCLA-113 |
| C*U | DL-30 | LA-119 | LA-111 | dCLA-119 |
| C*A | DL-30 | LA-118 | LA-111 | dCLA-118 |
| C*C | DL-30 | LA-123 | LA-111 | dCLA-123 |
| C*G | DL-30 | LA-112 | LA-111 | dCLA-112 |
| G*U | DL-30 | LA-108 | LA-111 | dCLA-108 |
| G*A | DL-30 | LA-114 | LA-111 | dCLA-114 |
| G*C | DL-30 | LA-110 | LA-111 | dCLA-110 |
| G*G | DL-30 | LA-124 | LA-111 | dCLA-124 |
| *U | DL-30 | LA-109 | DN-46 | dCLA-109 |
| *A | DL-30 | LA-113 | DN-46 | dCLA-113 |
| *C | DL-30 | LA-112 | DN-46 | dCLA-112 |
| *G | DL-30 | LA-123 | DN-46 | dCLA-123 |
| sU*sU | DL-30 | LA-122 | LA-111 | dCLA-122 |
| sC*sC | DL-30 | DN-24 | LA-111 | dCLA-124 |

##### 1.7.2 Measuring the kinetic parameters of modified guanosine imidazolium-bridged dinucleotides for Figures 6, S4.

The primer-template duplex was first annealed in a 40 µL solution containing 12.5 µM primer, 37.5 µM template, 50 mM NaCl, 50 mM Tris pH 8.0 and 1 mM EDTA, by heating at 94 °C for 2 minutes then cooling down to 12 °C at a rate of 0.1 °C/s. The 40 µL of annealing solution were diluted to a final volume of 100 µL in a solution containing 100 mM $MgCl_2$ and 200 mM Tris pH 8.0. A 5 µL aliquot of this solution was added to a 5 µL solution of the imidazolium bridged dinucleotide to start the reaction. 1 µL aliquots of the reaction were added to a quench solution containing 8M urea and 50 mM EDTA as well as 4.5 µM of the RNA complement of the template. Primer extension products were resolved by 20% (19:1) denaturing PAGE. The gel was scanned using a Typhoon 9410 scanner, and the bands were quantified using the ImageQuant TL software. The primers, template and complement sequence are listed below:

| Name | RNA or DNA | Name | Sequence (5′→ 3′) |
|---|---|---|---|
| Primer | RNA | DL-30 | /FAM/AGU GAG UAA CGG |
| Template | RNA | CG-1 | AAC CCC GUU ACU CAC U |
| Complement | RNA | CCG-1 | AGU GAG UAA CGG GGU U |

##### 1.7.3 Measuring the kinetic parameters of monomer-bridged-oligonucleotides for Figure 7, S5

The primer extension experiments were performed following the similar procedure as 1.7.1 but without downstream helper. The primers, templates, and complementary DNAs used in the primer extension assays are listed below:

| Monomer-bridged-oligo added | Primer | Template | Complimentary DNA |
|---|---|---|---|
| A*CG | DN-53 | DN-54 | dCDN-54 |
| A*CGC | DN-53 | DN-54 | dCDN-54 |
| A*CGCA | DN-53 | DN-54 | dCDN-54 |

### 2 Supplementary Figures and Tables

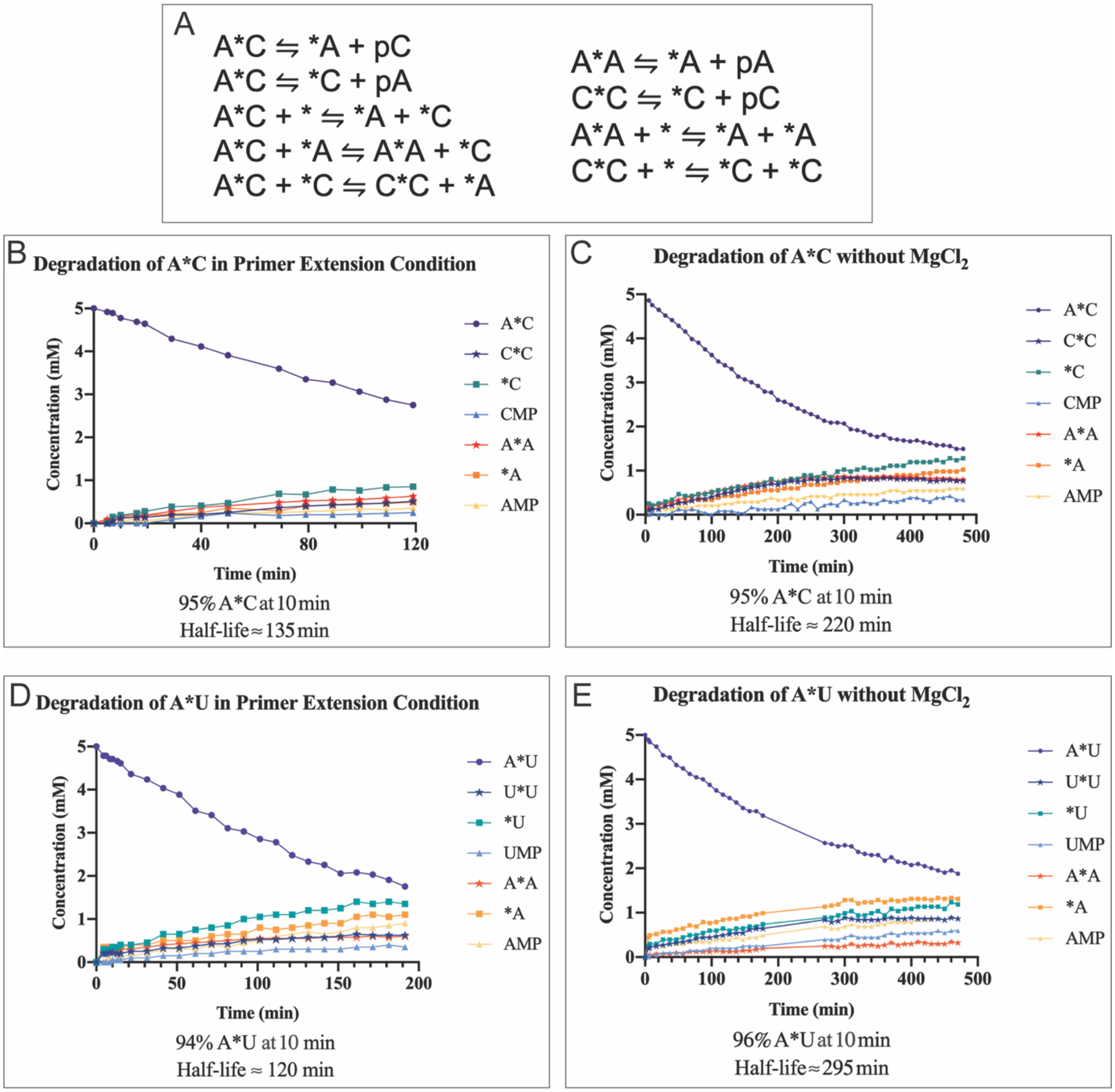


**Figure S1: Degradation of imidazolium-bridged dinucleotide intermediate measured by $^1$H NMR.** Degradation of A*C (top) and A*U (bottom) were measured in primer extension condition with magnesium anion (5 mM initial bridged dinucleotide intermediate, 100 mM $MgCl_2$, 200 mM d11-Tris pH 8.0) and without magnesium anion (5 mM initial bridged dinucleotide intermediate, 200 mM d11-Tris pH 8.0). (A) Possible reactions that can happen during the degradation of A*C (B) degradation of A*C under primer extension condition (C) degradation of A*C under primer extension condition without $MgCl_2$ (D) degradation of A*U under primer extension condition (E) degradation of A*U under primer extension condition without $MgCl_2$. The detailed experiment setup was described in section 1.6.

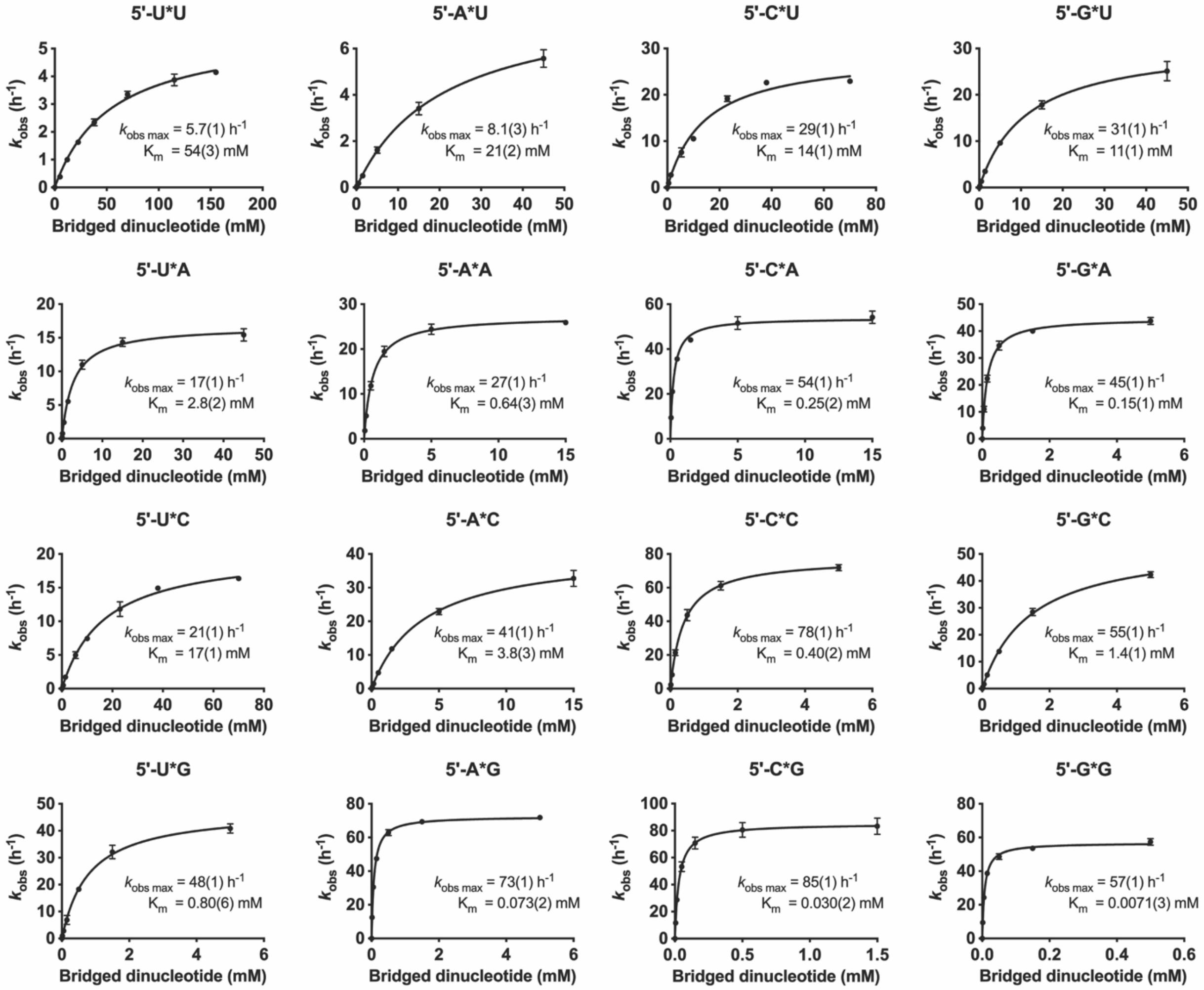


**Figure S2: Michaelis-Menten curves for primer extension with imidazolium-bridged dinucleotide intermediates.** The primer extension experiments were performed as described in section 1.7.1. The figures of the Michaelis-Menten curves are presented with the same order as in Figures 3 & 4. The $k_{obs\ max}$ and the $K_m$ for each curve are indicated to the right of the curve.

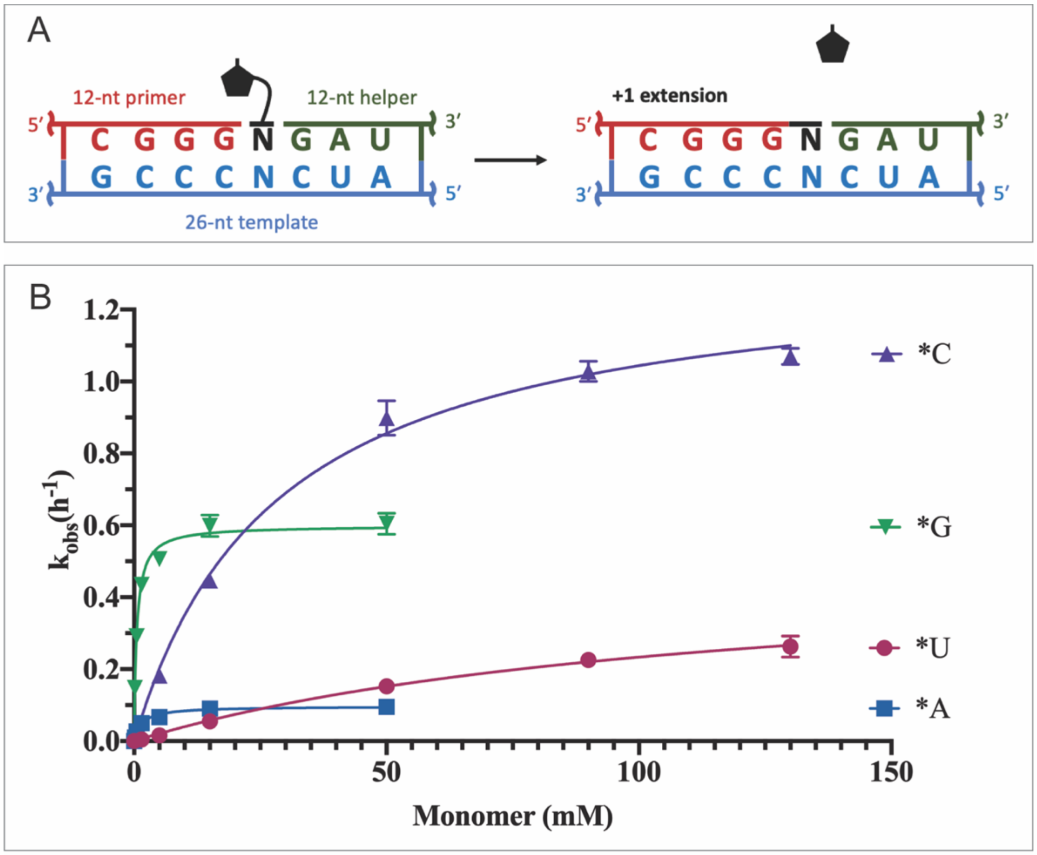


C

| | *C | *G | *U | *A |
|---|---|---|---|---|
| $k_{obs\ max}$ ($h^{-1}$) | 1.4(1) | 0.60(1) | 0.51(5) | 0.097(4) |
| $K_m$ (mM) | 29(2) | 0.54(4) | 120(20) | 1.5(3) |

**Figure S3: Primer extension with monomers in the sandwich system.** The primer extension experiments were performed as described in section 1.7.1. (A) Illustration of the primer extension experiment design. (B) Michaelis-Menten curves of the monomer primer extension. (C) Table of the $k_{obs\ max}$ and the $K_m$ of each monomer.

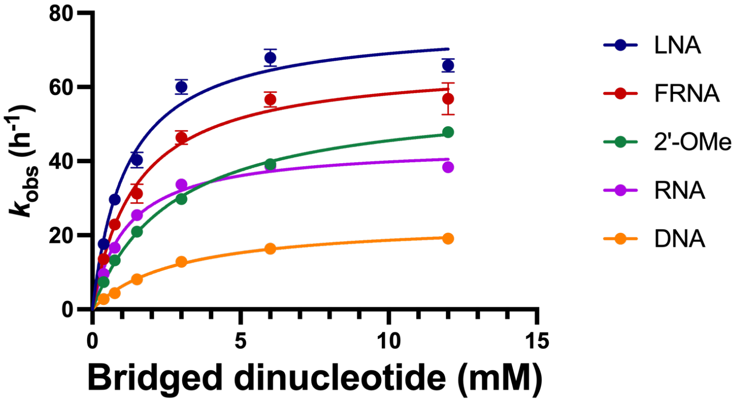


**Figure S4: Primer extension with different modified guanosine imidazolium-bridged dinucleotides.** The primer extension experiments were performed as described in section 1.7.2. The Michaelis-Menten curves are plotted together. The $k_{obs\ max}$ and the $K_m$ of each modified bridged dinucleotide is indicated in Figure 6.

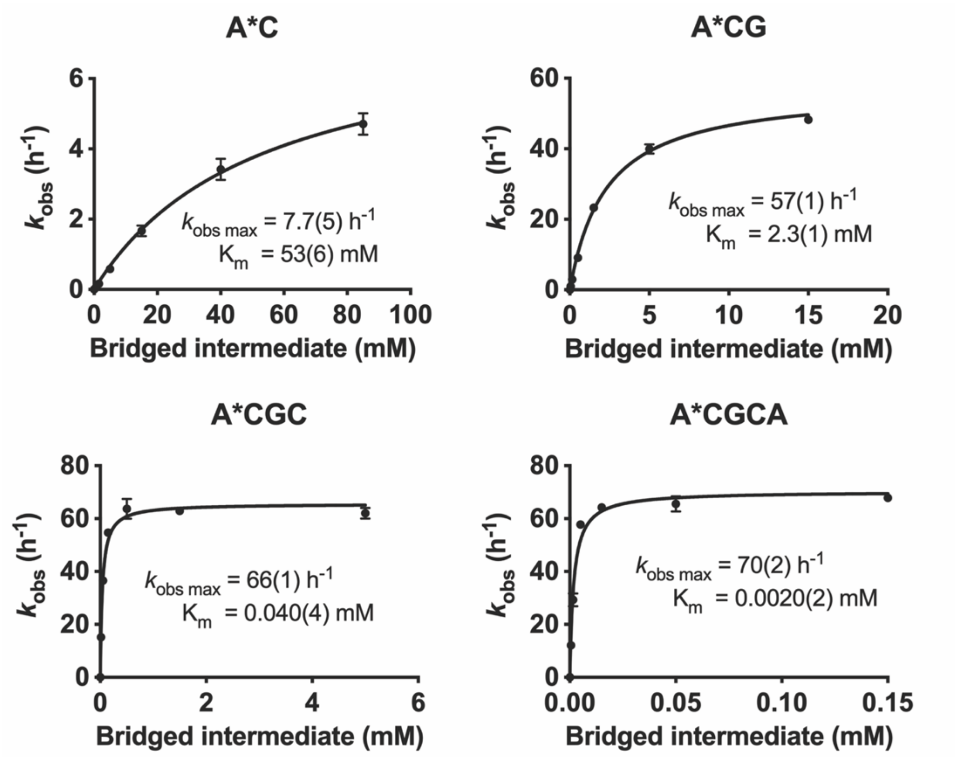


**Figure S5: Michaelis-Menten curves for primer extension with A*C, A*CG, A*CGC, and A*CGCA.** The primer extension experiments were performed as described in section 1.7.3. The $k_{obs\ max}$ and the $K_m$ for each curve are indicated at the right of the curve.

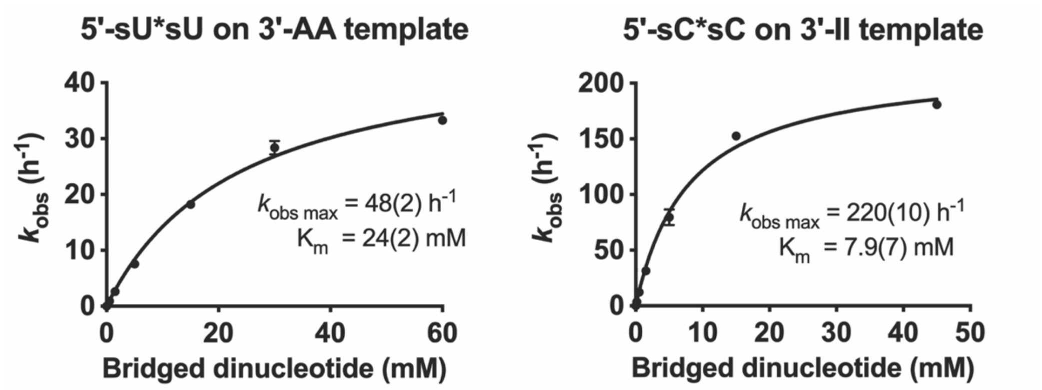


**Figure S6: Michaelis-Menten curves for primer extension with sU*sU and sC*sC.** The primer extension experiments were performed as described in section 1.7.1. The $k_{obs\ \mathrm{max}}$ and the $K_m$ for each curve are indicated at the right of the curve.

**Table S1: Sequences of oligonucleotides used in the study**

| Name | RNA or DNA | Sequence (5′→ 3′) |
|---|---|---|
| DL-30 | RNA | /FAM/AGU GAG UAA CGG |
| LA-111 | RNA | G AUG UCA GAU AU |
| DN-46 | RNA | GG AUG UCA GAU AU |
| LA-108 | RNA | AU AUC UGA CAU CAC CCG UUA CUC ACU |
| dCLA-108 | DNA | AGT GAG TAA CGG GTG ATG TCA GAT AT |
| LA-109 | RNA | AU AUC UGA CAU CCA CCG UUA CUC ACU |
| dCLA-109 | DNA | AGT GAG TAA CGG TGG ATG TCA GAT AT |
| LA-110 | RNA | AU AUC UGA CAU CGC CCG UUA CUC ACU |
| dCLA-110 | DNA | AGT GAG TAA CGG GCG ATG TCA GAT AT |
| LA-112 | RNA | AU AUC UGA CAU CCG CCG UUA CUC ACU |
| dCLA-112 | DNA | AGT GAG TAA CGG CGG ATG TCA GAT AT |
| LA-113 | RNA | AU AUC UGA CAU CCU CCG UUA CUC ACU |
| dCLA-113 | DNA | AGT GAG TAA CGG AGG ATG TCA GAT AT |
| LA-114 | RNA | AU AUC UGA CAU CUC CCG UUA CUC ACU |
| dCLA-114 | DNA | AGT GAG TAA CGG GAG ATG TCA GAT AT |
| LA-115 | RNA | AU AUC UGA CAU CUA CCG UUA CUC ACU |
| dCLA-115 | DNA | AGT GAG TAA CGG TAG ATG TCA GAT AT |
| LA-116 | RNA | AU AUC UGA CAU CAU CCG UUA CUC ACU |
| dCLA-116 | DNA | AGT GAG TAA CGG ATG ATG TCA GAT AT |
| LA-117 | RNA | AU AUC UGA CAU CGU CCG UUA CUC ACU |
| dCLA-117 | DNA | AGT GAG TAA CGG ACG ATG TCA GAT AT |
| LA-118 | RNA | AU AUC UGA CAU CUG CCG UUA CUC ACU |
| dCLA-118 | DNA | AGT GAG TAA CGG CAG ATG TCA GAT AT |
| LA-119 | RNA | AU AUC UGA CAU CAG CCG UUA CUC ACU |
| dCLA-119 | DNA | AGT GAG TAA CGG CTG ATG TCA GAT AT |

| | | |
|---|---|---|
| LA-120 | RNA | AU AUC UGA CAU CGA CCG UUA CUC ACU |
| dCLA-120 | DNA | AGT GAG TAA CGG TCG ATG TCA GAT AT |
| LA-121 | RNA | AU AUC UGA CAU CAA CCG UUA CUC ACU |
| dCLA-121 | DNA | AGT GAG TAA CGG TTG ATG TCA GAT AT |
| LA-122 | RNA | AU AUC UGA CAU CUU CCG UUA CUC ACU |
| dCLA-122 | DNA | AGT GAG TAA CGG AAG ATG TCA GAT AT |
| LA-123 | RNA | AU AUC UGA CAU CCC CCG UUA CUC ACU |
| dCLA-123 | DNA | AGT GAG TAA CGG GGG ATG TCA GAT AT |
| LA-124 | RNA | AU AUC UGA CAU CGG CCG UUA CUC ACU |
| dCLA-124 | DNA | AGT GAG TAA CGG CCG ATG TCA GAT AT |
| CG-1 | RNA | AAC CCC GUU ACU CAC U |
| CCG-1 | RNA | AGU GAG UAA CGG GGU U |
| DN-24 | RNA | AU AUC UGA CAU CII CCG UUA CUC ACU |
| DN-53 | RNA | /FAM/AGU GAG UAA CUC |
| DN-54 | RNA | UGCGU GAG UUA CUC ACU AAA |
| dCDN-54 | DNA | TTT AGT GAG TAA CTC ACGCA |
